# A Population Response Model of Ensemble Perception

**DOI:** 10.1101/2022.01.19.476871

**Authors:** Igor S. Utochkin, Jeunghwan Choi, Sang Chul Chong

## Abstract

Ensemble representations have been considered as one of the strategies that the visual system adopts to cope with its limited capacity. Thus, they include various statistical summaries such as mean, variance, and distributional properties and are formed over many stages of visual processing. The current study proposes a population coding model of ensemble perception to provide a theoretical and computational framework for these various facets of ensemble perception. The proposed model consists of a simple feature layer and a pooling layer. We assumed ensemble representations as population responses in the pooling layer and decoded various statistical properties from population responses. Our model successfully predicted averaging performance in orientation, size, color, and motion direction across different tasks. Furthermore, it predicted variance discrimination performance and the priming effects of feature distributions. Finally, it explained the well-known variance and set size effects and has a potential for explaining the adaptation and clustering effects.

---

Visual environment is composed of complex and dynamic objects, their relationships, and spatial layout. We stroll in a botanical garden and enjoy the spring blossom of flowers without bumping into a group of people passing by. Given the limited capacity of our visual system (Broadbent, 1958; Luck & Vogel, 1997; Palmer et al., 2011), how can we process the color of many flowers and the direction of people at the same time? Ensemble summaries provide one solution for the visual system to deal with this dilemma. The visual system does not have to represent the complex information in detail. Rather, it can summarize similar and redundant information as statistical properties to manage it efficiently within the limited capacity. Statistical summaries include average information, variability, and distributional properties. They are extracted and formed in multiple stages of visual processing and useful for many visual functions.

Average information is extracted from ensembles of similar objects more readily than their constituents, even if the exact average is not physically presented. When multiple items are present and over the capacity limit of the visual system, observers can judge the average information more accurately and rapidly than features of individual items (Ariely, 2001; Alvarez & Oliva, 2008; Haberman & Whitney, 2007; Parkes et al., 2001; Robitaille & Harris, 2011). Observers are likely to report having seen an item in a set, if it is similar to the average of the set (Ariely, 2001; Khayat & Hochstein, 2018, 2019). This tendency increases as a function of its distance to the mean. Moreover, the memory of individual items is biased toward their average (Brady & Alvarez, 2011; Corbett, 2017; Utochkin & Brady, 2020; Son et al., 2020), although they are not completely replaced with the average. Despite the bias, individual items are remembered more precisely if they are similar (Utochkin & Brady, 2020; Son et al., 2020). Therefore, the average information seems to be the best approximation of a set’s gist that is represented with relatively good precision and can compensate for less precisely represented constituents.

To represent a complex scene accurately, however, statistical summaries should include more than just a single representative value like the average. For example, we consider a variety of statistical properties, when we are shopping for a pack of cherry tomatoes in a farmer’s market. The average color of the cherry tomatoes indicates how ripe they are, and color variability indicates how homogeneous their ripeness is. We also consider their average size, size variability, and size distribution to choose the right pack. Accordingly, the visual system can accurately compute average color (Virtanen et al., 2020), color variability (Maule & Franklin, 2020), average size (Chong & Treisman, 2003), size variability (Solomon et al., 2011), and even richer information about the entire distribution of sizes (Kim & Chong, 2020) and colors (Chetverikov et al., 2017).

Moreover, the single mean for all items is not always a good representative of these items: for example, when yellowish flowers and greenish leaves are presented together on the same tree, their normal ranges of color variation are so different that there is of no use to calculate their common mean but it is better to get local means for each type of items. It appears that whether all items are considered a single group with one mean or as different groups with no single mean depends on the shape of a feature distribution (Treue et al., 2000; Utochkin, 2015). When the features are overall quite similar and smoothly distributed, they are likely to form a single-peak distribution with one mean as a good representative of the whole set. However, when there are a lot of highly dissimilar features and nothing in-between them, the distribution would have several peaks corresponding to more local groups. This suggests that the nuanced information about the feature distribution can be also an important aspect of ensemble representations.

Ensemble processing is involved with multiple stages of visual processing. Ensemble representations are accurately formed for low-level features such as orientation (Dakin & Watt, 1997), motion direction (Watamaniuk et al., 1989), speed (Watamaniuk & Duchon, 1992), color (Virtanen et al., 2020), brightness (Bauer, 2009), location (Alvarez & Oliva, 2008), and size (Ariely, 2001). They are also formed for high-level features such as facial expression (Haberman & Whitney, 2007), identity (Roberts et al., 2019), lifelikeness (Yamanashi Leib et al., 2016), and economic value (Yamanashi Leib et al., 2020). They are formed more accurately if included items are similar to each other (Ariely, 2001; Corbett et al., 2012; Dakin, 2001; Im & Halberda, 2013; Maule & Franklin, 2015; Sweeny et al., 2013; Solomon, 2010; Solomon et al., 2011; Utochkin & Tiurina, 2014) and if more items are included during averaging due to noise cancellation (Allik et al., 2013; Baek & Chong, 2020a; Brezis et al., 2018; Haberman & Whitney, 2010; Lee et al., 2016; Parkes et al., 2001; Robitaille & Harris, 2011; Solomon, 2010; Solomon et al., 2011)^1^.

Ensemble representations are ubiquitous and useful for many important visual functions. They can be used for learning and identifying categories (Khayat & Hochstein, 2019; Utochkin, 2015). The average information of a set can be considered as its prototype and its range can indicate a category boundary (Duffy et al., 2010; Khayat & Hochstein, 2019; Khayat et al., 2021). Previous ensemble representations are linked to current ensembles and individual representations to promote visual stability (Manassi et al., 2017). People use the distribution of distractor properties to find a target (Chetverikov et al., 2016; Treisman & Gormican, 1988; Utochkin & Yurevich, 2016). Stable ensemble representations facilitate visual search, even when they are not predictive of a target location (Corbett & Melcher, 2014). Ensemble representations are formed based only on a surface information automatically excluding contour information embedded in the surface (Cha & Chong, 2018), suggesting that ensemble representations can be used for figure/ground segregation. Finally, they are useful for texture perception (Cavanagh, 2001) like judging surface qualities such as glossiness (Motoyoshi et al., 2007). Ensemble information is an efficient way to represent information even about those objects whose conscious accessibility is severely diminished due to cluttering and crowding (Manassi & Whitney, 2018; Parkes et al., 2001) and, at the same time, ensemble representations of the surround can modulate the visibility of crowded objects (Tiurina et al., 2022).

Existing observer models of ensemble perception provide strong quantitative predictions that fit the performance of human observers well in some ensemble tasks. Most of the models account for averaging (Allik et al., 2013; Baek & Chong, 2020a; Parkes et al., 2001; Solomon et al., 2011) and only few of them go further to account for variability perception (e.g., Morgan et al., 2008; Solomon, 2010). These models are focused on quantifying observer inefficiencies in a set of parameters that could account for the patterns of performance found in experimental data. For example, the precision of average computation was shown to depend on limits imposed by the early noise involved with individual representations (Allik et al., 2013; Baek & Chong, 2020a; Im & Halberda, 2013; Parkes et al., 2001; Solomon et al., 2011), the number of items sampled from the entire set that can be sufficient to accomplish the observed error magnitude given the set distribution (Allik et al., 2013; Im & Halberda, 2013; Parkes et al., 2001; Solomon et al., 2011), late noise involved after sampling information about individual items and applied to readout ensemble summaries from this sample (Baek & Chong, 2020a; Parkes et al., 2001; Solomon et al., 2011), and distributed attention (Baek & Chong, 2020a). Whereas these models suggest accurate quantitative predictions about average and variance discrimination (Allik et al., 2013; Baek & Chong, 2020a; Parkes et al., 2001; Solomon, 2010; Solomon et al., 2011), their computational algorithms should not necessarily be taken for the mechanism of ensemble representation and their parameter estimates should not be taken to reflect independent representational states within that mechanism. Here are two examples of potential misconceptions that these models can entail if one attempts to directly map them onto ensemble mechanisms. First, model equations to estimate an observer’s efficiency directly implement regular statistical algorithms (e.g., the precision of average as a function of noise and effective set size follows the definition of the standard error of the mean). Therefore, it can be tempting to think that the computational mechanism behind ensemble representations also implements something similar to regular statistical calculations. For example, the arithmetic mean is defined as the sum of all sampled measurements divided by their number. However, it is not likely that the visual system does exactly that computation in the averaging task given that the judgments of sum of visible features are noisier than those of the average (Lee et al., 2016; Raidvee et al., 2021). The second example of a possible misconception concerns the idea of effective set size, a way to quantitatively describe the minimum amount of information an otherwise ideal observer should get from a display to reach the same level of efficiency as the real observer. It does not, however, literally refer to the number of individual items that the observer selects from the entire sets for further ensemble processing while ignoring the rest of the items (e.g., Myczek & Simons, 2008). This important point about the interpretation of effective set sizes in the ideal-observer models was explicitly made by Solomon (2021) in his recent review. Overall, the observer models provide a strong framework for quantitative description of behavioral patterns in various ensemble tasks, but they should not be necessarily considered to implement the mechanism of ensemble representation.

In the current paper, we propose a general, neurally plausible, and mechanistic model of ensemble representations across different feature domains. The model is based on population coding and spatial pooling, two very basic principles of organizing sensory systems of the brain. Many visual features are encoded as population responses, which implies that any particular feature value causes a specific pattern of responses (firing rates) in a group of neurons in accordance with their tuning to that feature (Brouwer & Heeger, 2009; Ester et al., 2009; Georgopoulos et al., 1986; Treue et al., 2000). Spatial pooling occurs from the hierarchical organization of visual processing that involves the progressively increasing size of receptive fields (areas of the visual field to which the neuron is responsive) from lower to higher brain regions (Boussaoud et al., 1991; Desimone & Schein, 1987; Ungerleider & Bell, 2011). Although previous studies have considered a population response model as a core mechanism of ensemble perception (Baek & Chong, 2020b; Chong & Treisman, 2003; Haberman & Whitney, 2012; Hochstein et al., 2018), they were descriptive outlines that have not been implemented as computational models, and thus they have not been tested against actual data. There are also a few examples of computational modeling of the global information representation with population coding model including motion or orientation integration (Dakin et al., 2005; Webb et al., 2007; 2010) or temporal number averaging (Brezis et al., 2018), but even these models were confined to explain averaging of a specific feature without generalization to other statistical properties or features. Their models employ multiple population responses of neurons selective to individual features or individual magnitudes (numbers) neurons, and a pooling layer to decode an average value from these responses. We propose an elaborated version of the population coding model that focuses on a key role of the pooling layer in ensemble perception. In our model, the pooled population response collected from multiple local responses to individual features is *the full-fledged neural representation of the ensemble* that preserves its rich statistical properties. This pooled response is sufficient to explain human performance in averaging, variance discrimination, and distribution-shape effects. In addition, our proposed model simulated well-known findings in the field of ensemble perception such as over-representation of mean information, the variance effect on averaging, outlier detection, and noise cancellation in large sets.

## Methods

### The architecture of the Core model

Our Core model of ensemble representation is an algorithm that simulates a pooled population response to a stimulus set of individual features; we operationalize this population response as a neural code of the ensemble. The Core model can be presented as a two-layer neural network, with the bottom layer (*L1*) representing individual items in small spatially separated receptive fields and the top layer (*L2*) pooling signals from the bottom layer within a single receptive field (Figure 1). Units of these two layers are feature-selective neurons tuned to fire at a certain rate depending on stimuli. Connections between the layers are feedforward and each bottom-layer unit is connected to each top-layer unit with a certain synaptic weight. The distributions of synaptic weights can be conceptualized as tuning functions of the pooling top-layer units. In a fully biologically plausible system, stimuli would cause population response in both layers, so that a number of neurons is differentially activated when a given feature is presented. However, for simplicity, we conceptualize responses of the bottom layer as singular feature values corresponding to the actual stimulus value (corrupted by some amount of early noise). We can make this simplification assuming that population responses of a lower layer are already included in measurable outputs of the top-level units, which are used to plot tuning curves. For example, the tuning curve of monkey V4 can be built based on direct measurements from V4 neurons, however, the firing rate of each individual V4 neuron in fact results from the population activity of multiple lower-level neurons. Researchers usually omit the hidden population activity of lower layers, so the tuning curve of the measured neuron usually is plotted directly as a function of stimulus. Therefore, we omit the population responses of the bottom-layer units because we use the same logic.

**Figure 1.**
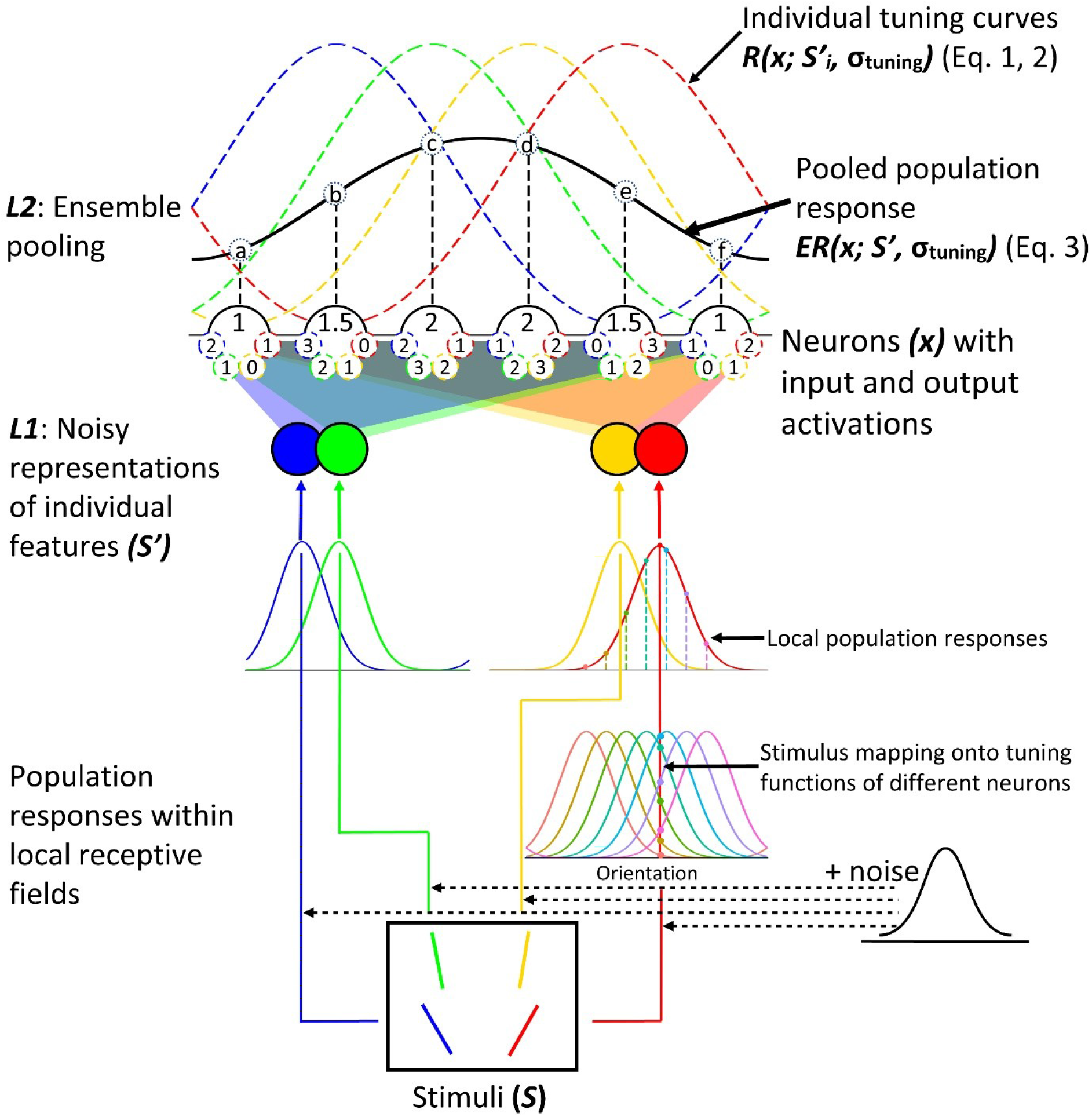
The Core model of pooling and population coding explaining the formation of a neural representation of ensemble. *Note.* The bottom layer (*L1*) of the neural network (filled color squares) depicts simplified outputs of local population responses. Each unit (blue, green, yellow, and red solid curves) represents a noisy population code of a single item (*S’_i_*). In the bottom right corner, we illustrate how a red oriented bar produces such a population code based on tuning functions of individual neurons from the population. The top layer (*L2*) is a population of selectively tuned neurons (shown ordered horizontally by their orientation preference *x*) pooling local signals from the bottom-layer units (the values of input signals are shown inside small “synapses” around each *L2* unit and the values of the pooled responses are shown inside the *L2* units themselves). The distribution of synaptic weights corresponds to component tuning curves (*R*(*x; S’_i_*, *σ*_tuning_)) of the top-layer neurons (blue, green, yellow, and red dotted curves, respective to the colors of the stimuli and their *L1* representations). Outputs of the top-layer neurons are the sums of contributing bottom-layer inputs with their synaptic weights. The shape of the population response (neural representation of an ensemble termed *ER*(*x; S’_i_*, *σ*_tuning_)) following from the distribution of top-layer outputs is depicted as a black curve and each point (black numbers: a, b, c, d, e, and f) on the curve is computed based on the average (neural normalization) of the top-layer neuron outputs.

The wiring scheme of connections between the bottom and top layers is critical to understand the machinery of the pooling mechanism that creates the neural representation of the ensemble. This wiring scheme defines feature preferences (tuning curves) of the top-layer neurons. In Figure 1, it is shown as a set of synaptic weights (we illustrate them with different numbers). In our example, the maximum weight is 3 that shows perfect matching between the activated bottom-level and top-level units; weights 2, 1, and 0 show response attenuation as a function of decreasing similarity between the preferred and actually presented values (Note that the synaptic weights as well as the shape of the tuning function in Figure 1 are arbitrary and used only for illustration purposes). As the top-layer units are pooling, we can present the outcome activation of each such neuron as the average of synaptic weights produced by each active bottom-layer unit. Note that signal averaging is a neurally plausible way of pooling local responses to different competing visual features by neurons with large receptive fields when no stimulus is selectively attended (Desimone & Duncan, 1995; Kastner et al., 1998). The distribution of individual top-layer outcomes defines the shape of the pooled population code corresponding to an ensemble representation.

### Computational modeling

Computational modeling of ensemble perception was implemented in codes written in R language. We have run a set of simulations of various studies from the ensemble literature using different tasks, feature dimensions, and feature distributions. Each simulation was based on the Core model producing a pooled population response and also included a specific part depending on a simulated paradigm. The specific part in most cases included reading-out task-relevant output parameters of the population code, response generation based on these parameters, and performance estimation (e.g., various accuracy metrics – percent correct, just noticeable difference, and *d*’). Model performance was then tested for correlation against the data from the corresponding studies in order to estimate how well the model predicts the observed patterns. The only exception was the feature distribution learning (FDL) paradigm (Chetverikov et al., 2016, 2017) where population codes produced by the Core model were directly compared against the behavioral data, because the paradigm does not imply any explicit report about ensemble per se (see detailed explanation in Modeling feature distribution learning section of Methods).

For user convenience, all of our simulations were implemented as a set of functions using only stimulus settings as arguments. These functions are collected in the R package, *‘EnsPopulationModel’* that can be found at https://osf.io/9yxv5/. This study was not preregistered.

#### Computational implementation of the Core model

The Core model was computationally implemented as a function (*‘popcode’* function) using a vector of feature values of a stimulus set and a feature space specification as arguments. The output of the function is the distribution of spike rates among a set of feature selective neurons *x* ordered by their feature preference whose preference range and granularity are predefined in a separate numeric vector.

The bottom layer (*L1,* Figure 1) of the Core model takes each individual value from the input stimulus set *S* ∈ {*S*_1_, *S*_2_, … *S*_n_} and corrupts it by the early encoding noise drawn from a normal distribution with a standard deviation σ*_E_* and truncated by 2 standard deviations. It resulted in a set termed *S’* such that *S’ ∼ S + N(*0, σ*_E_)*. The *S’* models the noisy responses to individual items in neural populations with small receptive fields (roughly, V1-like neurons). The *S’* is passed to the top layer (*L2,* Figure 1).

The top layer of the model pools the noisy signals of the bottom level. For each individual bottom-layer signal, a population response is produced in all top-layer neurons in accordance with synaptic weights approximated by a normal (for linear dimensions) or a wrapped normal (for circular or semi-circular dimensions such as orientation, color hue, and motion direction) probability density function.

For linear dimensions:

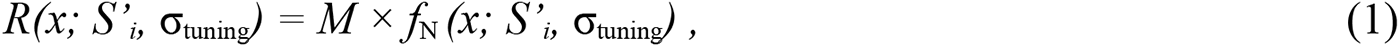

where *R(x; S’_i_, σ_tuning_)* are the responses of neurons from a population of neurons *x* ordered by their stimulus preference to a bottom-level individual signal *S’_i_* from vector *S’,* σ_tuning_ is a standard deviation of the Gaussian tuning curve of the top-layer neurons, and *M =* 30*/* max[*f*_N_*(x; S’_i_, σ*_tuning_)] is the spike rate of a neuron at the peak of its feature preference. Since *M* is a constant scaling factor, it can be set arbitrarily to fit neurally plausible values (we set it at 30 Hz in our simulations), but it does not affect the shape of the population response.

For circular and semicircular dimensions:

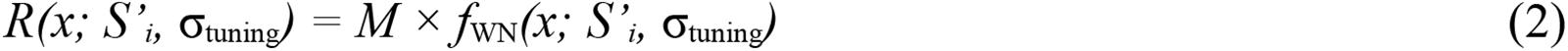

Note that for the circular and semicircular dimensions, *x*, *S’_i_*, and σ_tuning_ are angular variables entered in degree units. To map these values onto the circular dimension, they were converted to radians as *θ°*×π/180, where *θ°* is the degree value of an argument variable. To map these values onto the semicircular dimension, we used the conversion formula of *θ°*×π/90.

The *R(x; S’_i_,* σ_tuning_) along the whole population *x* can be interpreted as a population response to an individual stimulus representation *S’_i_* if this stimulus is presented alone. In terms of the joint (ensemble) population response, *R(x; S’_i_,* σ_tuning_) is its individual component. Therefore, the contributions of all individual components to the activation of the top-level response will be defined by a matrix C = ‖*R(x_j_; S’_i_,* σ_tuning_)‖, where *R(x_j_; S’_i_,* σ_tuning_) is the *i*-th component activation of the *j*-th top-layer neuron.

The pooled ensemble population response *ER(x; S’,* σ_tuning_) is accomplished by averaging all *n* components for each of the *m* top-layer neurons:

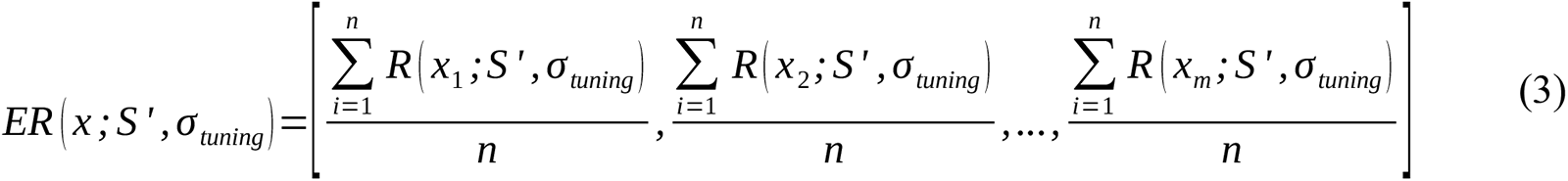

Figure 2a shows four illustrative examples of the pooled population code resulting from our Core model as a function of the feature distribution in a stimulus set with different orientations. These four examples demonstrate how the population response takes a shape isomorphous to the feature distribution and thus conveys its critical properties. Green, red, and blue curves in Figure 2a show population responses to smoothly distributed orientations with different means and variability. Most notably, middle neurons accumulate more local signals, as they are activated by features from both sides of the synaptic circuit, whereas neurons at the edges are activated only from one side. The overall distribution does, therefore, have a single peak around the mean prevailing over the activity of other neurons. The comparison between red and blue curves in Figure 2a illustrates how an ensemble representation depends on feature variability in a display. When a greater range of features is involved, activation spreads to a larger population of pooling units, so the bandwidth of an active population gets broader as well. Finally, the black curve in Figure 2a shows a population response to a display with highly dissimilar features. Here, clusters of similar features (e.g., all steep lines) contribute to the activation of local subpopulation of pooling neurons but the absence of features in the middle of the physical feature distribution prevents the formation of a single peak in the middle of the population code. Instead, two local peaks are formed that can correlate with the perception of two highly distinct groups of features rather than a single set (Treue et al., 2000; Utochkin, 2015).

**Figure 2.**
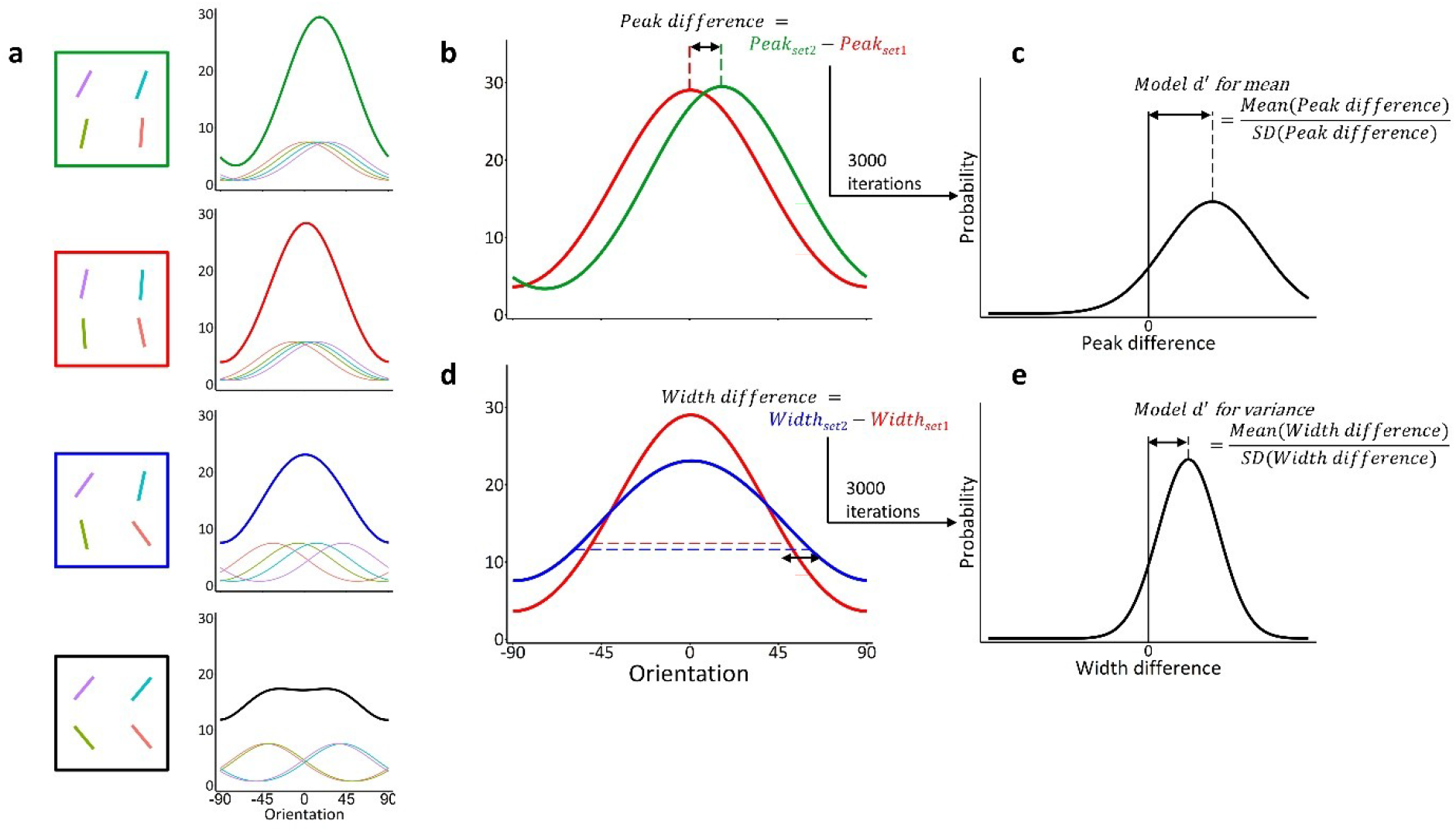
Population responses of the pooling layer (V4-like neurons) to 4-item ensembles of oriented lines. *Note.* (a) narrow uniform distribution (green & red), broad uniform distribution (blue), and two-peak distribution with extremely spaced orientation values (black). The pooled population response is shown by the thick lines and corresponds to the ensemble neural representation; the thin colored lines depict component population responses to individual stimuli (amplitudes divided by set size due to averaging). The peaks of individual curves are jittered because of the early noise. Our model computes a peak difference from two population responses for 3000 times (b) to simulate a *Model d***’** for mean discrimination (c). Similarly, it computes a width difference from two population responses for 3000 times (d) to simulate a *Model d***’** for variance discrimination (e).

#### Output model parameters

The resulting population code *ER(x)* from equation 3 has a set of output parameters that can have straightforward interpretation in terms of ensemble summary statistics that observers might estimate in a stimulus. The *peak location, P* can be naturally interpreted as the most likely estimate of an ensemble mean. It is defined as the feature preference of neuron having the maximum firing rate *ER(x)* in the population *x*:

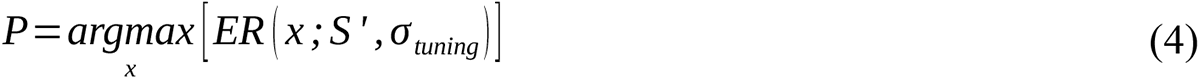

The bandwidth of the population activity, calculated as the standard deviation of the involved pooling neurons weighted by their outcome firing rates, can be used as an estimate of ensemble variability. We term this parameter a *population width (W)*. For linear feature dimensions, the *W* was calculated as a weighted standard deviation and specified as *W_linear_*(that in our code was implemented by the function ‘w.sd’ from the R package ‘weighted.desc.stat’: https://cran.r-project.org/web/packages/Weighted.Desc.Stat/index.html):

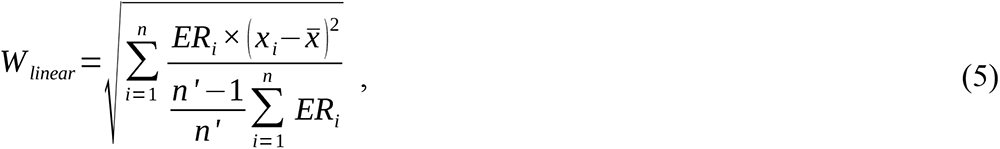

where *x_i_* is the feature preference of the *i*-th neuron, *ER_i_ = ER(x_i_; S’,* σ_tuning_) is the output firing rate of the *i*-th neuron, 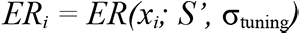 is the weighted average preferred feature of the population, *n* is the number of neurons in the population, *n’* is the number of non-zero output firing rates.

For circular and semi-circular feature dimensions, the *W* was defined as a weighted circular deviation. The weighted circular deviation is based on the length of the mean resultant vector *r*:

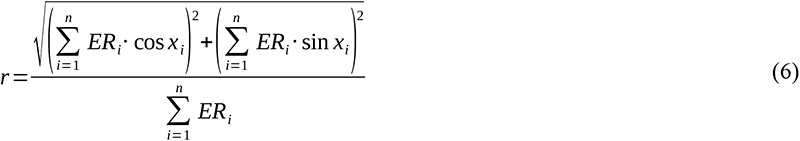

Then, the *W_circular_* is calculated using the equation for the circular deviation (Berens, 2009):

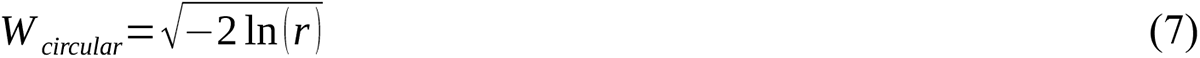

In our computational model package, two functions are used to calculate the peak location *P* and the population width *W*. The first function, *‘decode’* has feature values of a stimulus set *S* as an argument and returns the *P*, the *W*, as well as the real mean feature and real standard deviation of the set. The second function, *‘decode.direct’* has a pooled population response (the output of the Core model, *ER*(*x; S’,* σ_tuning_)) as an input and returns the *P* and the *W*.

#### Modeling mean and variance discrimination in 2-AFC tasks

In each trial of a typical two-alternative forced choice task (2-AFC) or two-interval forced choice task (2-IFC), observers are shown two sets of items either in two distinct locations simultaneously or in the same location serially. The observers are instructed to determine the direction of difference between the sets along a target dimension. For example, in the orientation averaging task (Figure 3a), observers are asked to answer whether set 2 is tilted more clockwise or counterclockwise than set 1 on average (e.g., Solomon, 2010) or vertical (e.g., Dakin, 2001; Yashiro et al., 2020). In a variance discrimination task, observers are instructed to answer whether set 2 has greater or smaller feature variation than set 1. Regardless of the particular task, the physical difference between the sets is systematically manipulated across trials that allows to estimate the discrimination threshold (also referred to as the just noticeable difference, JND) via the proportion of “set 2 greater” against “set 1 greater” answers as a function of the physical difference (psychometric function).

**Figure 3.**
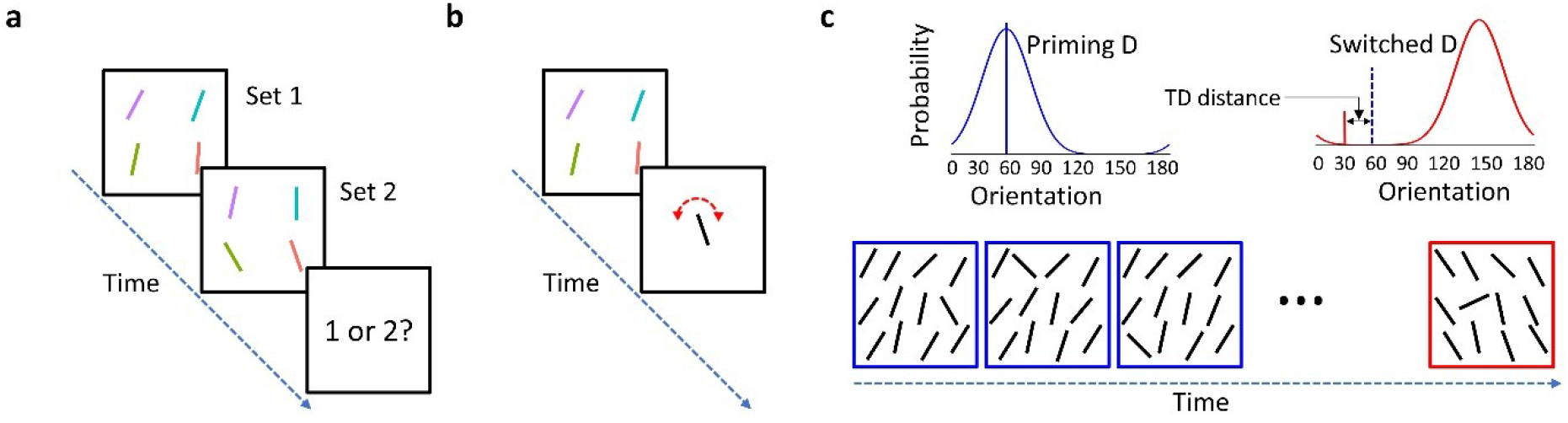
Various research methods used in the ensemble coding literature. *Note.* (a) In a two-interval forced choice task, observers are asked to judge whether Set 2 is more clockwise or counterclockwise (or greater or smaller for linear feature dimensions) than Set 1 on average. (b) In an adjustment task, observers are typically shown a set of oriented bars and are asked to set a subsequently presented adjustable probe as close as possible to their mean orientation. (c) In feature distribution learning experiments, observers are first exposed to a streak of odd-one-out visual search trials (shown as blue framed displays) based on a consistent distractor distribution (blue priming distribution) and a switch trial (shown as a red framed display) based on a switched distribution with a shifted mean orientation (red distribution) and a target (a red bar) drawn from the feature range that can fall within the range of the priming distribution. A dotted blue bar indicates the mean of the priming distribution.

##### Simulating observer behavior from pooled population responses

In each run of our 2-AFC simulations, we generated two vectors of values corresponding to individual items in sets 1 and 2. Set sizes and distributional properties of sets were generated in accordance with methods described in the relevant published studies. The absolute physical differences between sets 1 and 2 ranged from 0 to a magnitude at least twice as large as the highest JND value reported in a referred study or set of studies. These two sets of stimulus values were the model input. A random number drawn from a Gaussian distribution (mean = 0, different *SD*s depending on feature dimension) was then added to each individual value to simulate stimulus corruption by the early noise (σ*_Ε_*) in layer 1. Pooled population responses, as described in our Core model, were then created for each of the sets. The tuning curves of pooling neurons were Gaussian functions with a *SD* depending on feature dimension.

When the population responses for sets 1 and 2 were created their output parameters could be read out to produce an answer on a given trial. For the averaging tasks, a difference between peak locations of population response 2 and population response 1 was read-out (Figure 2b), whereas for the variance discrimination tasks, a difference between population widths between these two population responses was read out (Figure 2d). If the difference between the peaks or between the widths was positive, then the model observer answered “set 2 greater”; if the difference was negative, the model observer answered “set 2 smaller”; if the difference was zero the model observer chose one of these responses randomly. After running 3,000 simulations on each condition (each combination of feature distribution, set size, and physical difference between sets 1 and 2), the probability of “set 2 greater” was calculated (Figures 2c and 2e). The distribution of these probabilities as a function of the physical difference (psychometric function) was fit as a cumulative Gaussian distribution and the JND was determined using the same threshold as in the reference study.

##### Signal-detection parameters from the model and behavior

The peak location and the bandwidth parameters of the population code normalized by corresponding variability measures make psychophysically meaningful proxies of sensitivity to ensemble summary statistics. For any given stimulus difference either along the average feature or along feature variance, percent correct answers can be converted to a behavioral discriminability index, *d’*, which reflects a signal-to-noise ratio, the mean distance between the representations of the two sets in the feature space normalized by some internal noise (Macmillan & Creelman, 2005). This internal noise corrupts representations of the sets (or of the set and the comparison item) and makes them confused on a certain proportion of trials. Our model can produce an analog of such *d’* for a 2-AFC task. For each of two sets (or for a set and an item), two population codes are generated and the difference between their relevant output parameters – difference between peak locations for the average comparison task (Figure 2b), difference between bandwidths for the variance comparison task (Figure 2d) – is calculated. Depending on whether the resultant difference is positive or negative, a decision is made as to which one of the sets has a greater mean or variance. Since the early noise makes peak and bandwidth parameters fluctuate from trial to trial, the differences between the representations will also fluctuate which will provide a bell-shaped distribution of representational differences across trials (Figures 2c and 2e). The mean of this difference distribution can be conceptualized as the neural difference signal between the population responses. The greater the mean is shifted away from 0, the stronger the signal. The *SD* of the difference distribution can be conceptualized as the amount of discrimination noise. The mean difference normalized by the SD (*M_diff_/SD_diff_*) is the signal-to-noise ratio, or the model’s *population d’* (Figures 2c and 2e). Note that the model’s population *d’* is not supposed to equal the behavioral *d’* that can include other sources of uncertainty (e.g., related to decision) not considered by our model. Yet, we consider the introduction of the population *d’* useful as a sensible and measurable concept that provides simple mapping of the population code model onto psychophysical measurements in behavioral experiments.

The behavioral *d’* for a given condition can be obtained from the observed JND assuming that the psychometric function, from which this JND is calculated, is a cumulative normal distribution. Since the JND is the stimulus difference that provides a known probability of correct discrimination, this probability can be *z*-transformed and the *SD* of the psychometric function can be found. Based on the *SD*, a *z*-score for any physical difference between two sets can be calculated and used to find the 2-AFC *d’* (see the formula in Stanislaw & Todorow, 1999). The distributions of behavioral and model *d’*s across various stimulus conditions and in a fixed physical difference can be further used to estimate their correlation that would show the quality of model predictions regarding behavior.

#### Modeling averaging in the method of adjustment

In the method of adjustment (Figure 3b), observers are typically shown a single sample stimulus (e.g., a set of items) and are asked to set a subsequently presented adjustable probe as close as possible to the target feature of the sample (e.g., mean orientation). Since the probe can be adjusted in an almost continuous fashion, the data are usually presented as a continuous error distribution across trials. The mean of this distribution is taken for the point of subjective equality and its standard deviation is taken as discrimination precision measure (e.g., a JND). Since there is only one set presented, we simulate our model responses in the adjustment method by simply picking a corresponding output parameter (e.g., peak location for averaging) of a simulated pooled response. The average output parameter across runs is taken for the point of subjective equality and the standard deviation is taken for precision (not including late noise factors).

#### Modeling feature distribution learning

##### Behavioral data

In feature distribution learning (FDL) experiments (Figure 3c), observers are first exposed to a streak of trials with a consistent distractor distribution and an odd-one-out target (for example, a line with a highly distinctive orientation or a diamond with a highly distinctive hue – Chetverikov et al., 2016, 2017). After priming an observer with such a streak of trials, a switch trial follows where the distractor distribution dramatically shifts its mean feature (and can also change its shape) and the target feature also takes a new value that can fall within the range of the former distractor distribution (hereinafter, the *priming distribution*), as well as beyond this range. Observers’ task was always to find the target as fast as possible and to respond as to which part of the array it is located. Following the principal logic of the FDL paradigm, our analysis was focused on reaction times (RT) in the switch trials where priming effects of the previously learned distractor distribution are tested. The distribution of RT’s in the switch trials as a function of the target feature distance from the middle-range feature of the former distractor distribution (hereinafter, TD distance) was our main subject of prediction. We also applied local polynomial regression fits of RT × TD distance functions (same as in Chetverikov et al., 2016, 2017). Correlation coefficients between the RT × TD distance data (binned by 5 degrees) and the regression fits were considered to reflect the “ceiling” quality of the fits that might be reached with a smooth continuous function based only on the data. The correlations between the data and model predictions are further compared with these ceilings.

##### Physical priming distributions

In order to see how much the RT distributions are predicted by the distributions of physically presented features, we directly used probability density functions as in Chetverikov et al. (2016)’s experiments that we chose for modeling: Gaussian distributions with SD = 10 degrees and SD = 15 degrees, a uniform distribution covering from -30 to 30 degrees (all three distributions tested in Chetverikov et al. (2016)’s Experiment 2), and two skewed, triangular distributions ranging from -30 to 30 degrees, either right-skewed (peak = – 25 degrees), or left-skewed (peak = 25 degrees) (these two shapes were tested in Chetverikov et al. (2016)’s Experiment 4). The probability densities based on these theoretical distributions were correlated with the RT × TD distance data in the corresponding priming distribution conditions to estimate how well the RT distributions are predicted by the real feature distributions.

##### Population model distributions

To estimate how much the RT distributions primed by a certain distractor distribution can be predicted by the ensemble population response, we estimated the correlation between the RT distributions and the average spike rate distributions provided by our core model for corresponding priming distributions. We had 3,000 runs for each type of the priming distribution (as listed in the previous paragraph). In each run, we randomly drew 35 feature values from a defined distribution, each feature value corresponded to one individual distractor from Chetverikov et al.’s (2016) standard displays (they always presented an array of 36 elements, one of them reserved for the target and the rest 35 for distractors). In accordance with our core model, these 35 individual features were corrupted by the early noise and then pooled by neurons with the default tuning parameters. The output spike rate distributions were then averaged across runs to get the model prediction of the ensemble population response.

Note that for visualization purposes we normalized RT’s, probability densities of features in physical priming distributions, and spike rates in model’s population responses to fit these functions in the same scale from 0 (minimum value) to 1 (maximum value). Note that this normalization does not affect any correlations between the distributions, as it only changes the scale but not relations between the points.

### Modeling results

We applied our ‘pooling and population coding’ model to different paradigms and feature dimensions used in previously published studies (Table 1) to see whether our model can account for the variety of data accumulated in the field. To remind, the model focuses on the relatively early representational mechanism of ensemble perception that is limited by the early input noise and by tuning characteristics of pooling neurons. The model does not include late sources of uncertainty and noise (such as readout error, decision error, or lapses). Therefore, our model naturally does not predict behavioral data exactly. We demonstrate instead that, although model predictions and data may be differently scaled, the model nicely captures basic patterns and trends observed in the data (including some interesting non-linearities and non-monotonicity, such as the “dipper function” in variance perception – see details below). The principal significance of these demonstrations is that a broad range of effects can be captured by exactly the same simple mechanism without introducing any additional assumptions.

**Table 1.**
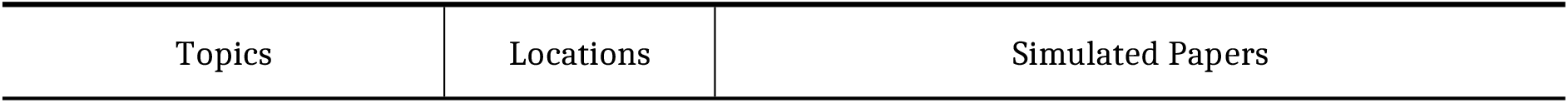
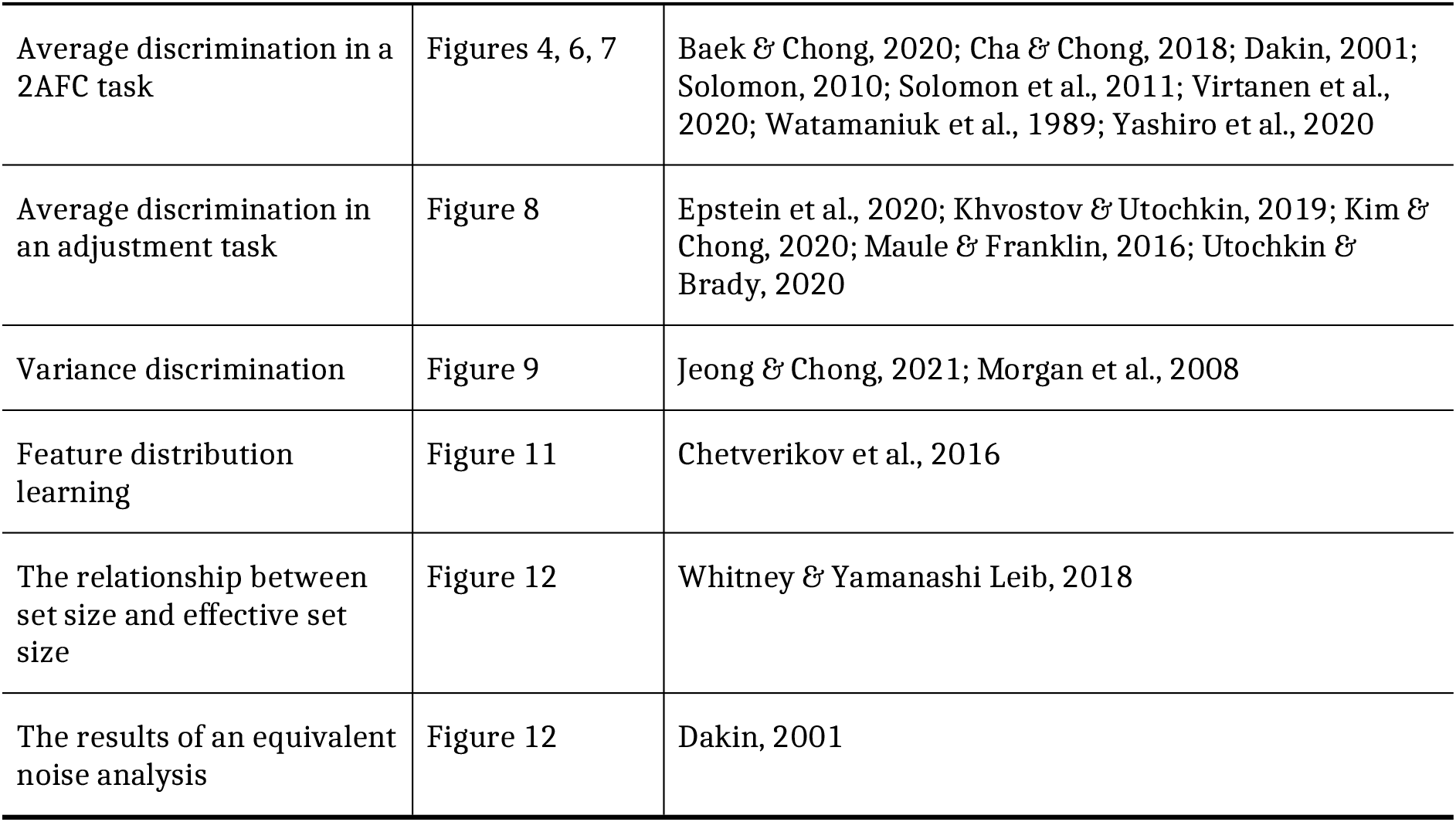
The simulation list of published studies reported in the present study.

| Topics | Locations | Simulated Papers |
| --- | --- | --- |
| Average discrimination in a 2AFC task | Figures 4, 6, 7 | Baek & Chong, 2020; Cha & Chong, 2018; Dakin, 2001; Solomon, 2010; Solomon et al., 2011; Virtanen et al., 2020; Watamaniuk et al., 1989; Yashiro et al., 2020 |
| Average discrimination in an adjustment task | Figure 8 | Epstein et al., 2020; Khvostov & Utochkin, 2019; Kim & Chong, 2020; Maule & Franklin, 2016; Utochkin & Brady, 2020 |
| Variance discrimination | Figure 9 | Jeong & Chong, 2021; Morgan et al., 2008 |
| Feature distribution learning | Figure 11 | Chetverikov et al., 2016 |
| The relationship between set size and effective set size | Figure 12 | Whitney & Yamanashi Leib, 2018 |
| The results of an equivalent noise analysis | Figure 12 | Dakin, 2001 |

#### Average discrimination in a 2-AFC task

2-AFC discrimination of mean orientations has been employed in a number of studies (Cha & Chong, 2018; Dakin, 2001; Dakin & Watt, 1997; Solomon, 2010; Rosenholtz, 2000). In some of these works, researchers have parametrically manipulated set size (the number of elements in a set) and the physical variance of sets (e.g., Dakin, 2001; Solomon, 2010). They found the variance and set size effects. The variance effect shows that the mean discrimination threshold, as calculated from the slope of a psychometric function (the proportion of answering that one stimulus is greater than another as a function of the physical difference between them), is plateaued at the beginning (when stimulus variances are up to 3-4 degrees) and then steadily grows as a function of variance (Figure 4a). The variance effect is robust not only in an orientation domain but also in other domains such as color (Maule & Franklin, 2015) and size (Im & Halberda, 2013; Solomon et al., 2011). The set size effect indicates that mean discrimination threshold generally improves with increasing set size (Figures 4a). Such a benefit from larger set sizes can be also found in some of the size averaging experiments (e.g., Allik et al., 2013; Baek & Chong, 2020a; Chong et al., 2008; Robitaille & Harris, 2011), although other studies failed to report it (Ariely, 2001; Marchant et al., 2013). Existing accounts in the computational models of inefficient observers usually capture this effect by allowing some internal parameters change such as effective set size (Allik et al., 2013; Dakin, 2001; Dakin & Watt, 1997; Im & Halberda, 2013; Maule & Franklin, 2016; Solomon, 2010; Whitney & Yamanashi Leib, 2018), the variable amount of early noise (Alvarez, 2011), or attention (Baek & Chong, 2020a).

**Figure 4.**
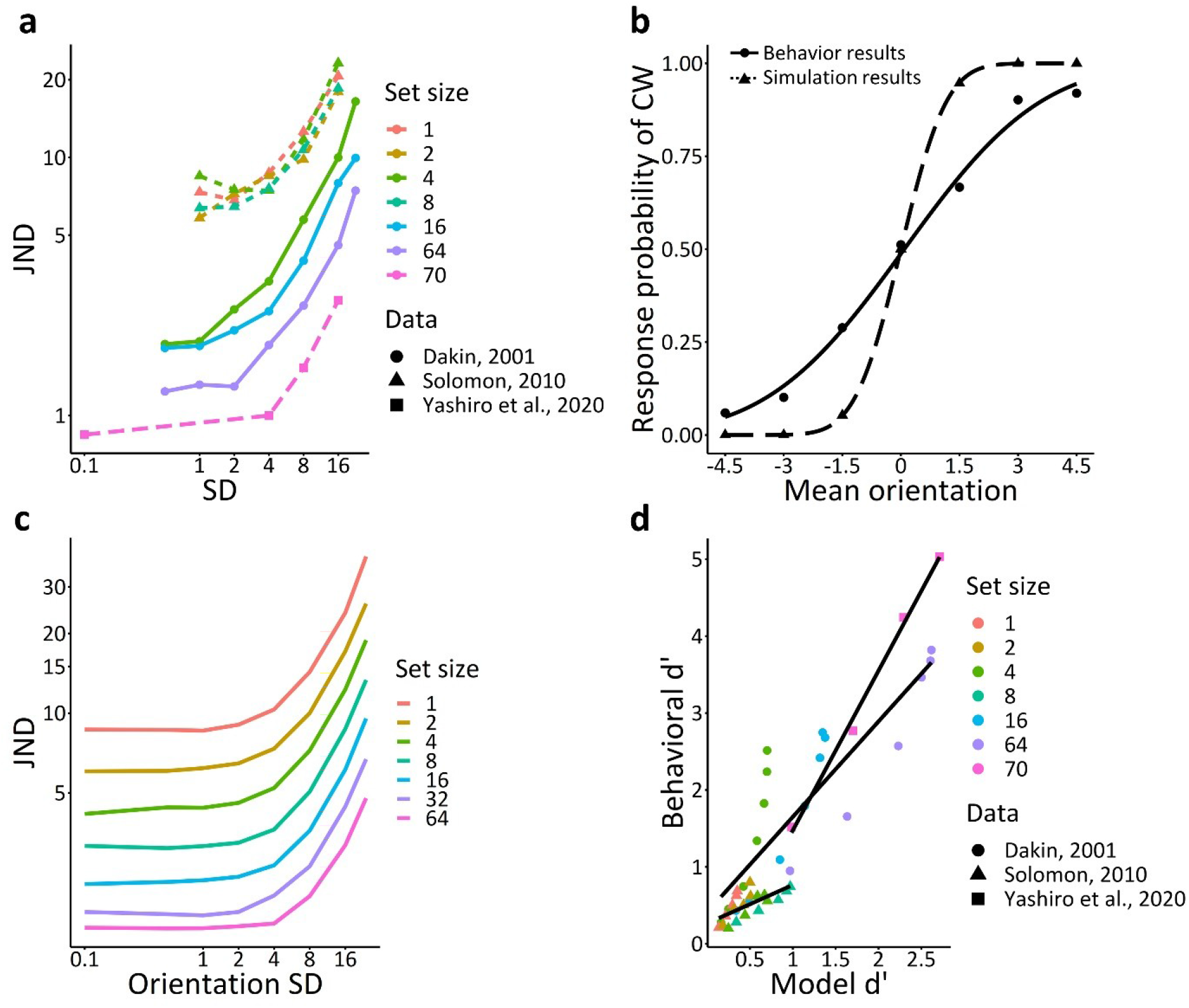
The variance and set size effects on average discrimination in a 2AFC paradigm. *Note.* (a) Previous studies have found that mean orientation discrimination performance decreased with increasing variance and decreasing set size (Dakin 2001; Solomon, 2010; Yashiro et al., 2020). (b) Psychometric functions of orientation discrimination. The solid line indicates the results from Cha & Chong (2018) and the dotted line is the results of the model simulation. (c) Model JNDs are plotted against SD and set size. They capture the qualitative patterns of behavioral results presented in (a) well. (d) The correlation results between behavioral *d*’ from previous studies (Dakin 2001; Solomon, 2010; Yashiro et al., 2020) and model *d*’.

##### Orientation

Our model explains both the plateau-and-growth variance effect and the facilitating set size effect^2^ on the discrimination of average orientations. Noteworthy, the model does it parsimoniously, without the need to fit free parameters to every situation. We used a fixed set of parameters based on the previous research of orientation averaging and orientation population coding. Namely, the width (SD) of the Gaussian tuning curve was set at 37 degrees (McAdams & Maunsell, 1999) and the SD of the Gaussian early noise was set at 7 degrees (Dakin, 2001; Solomon, 2010). Figure 5 shows how our model simulates mean orientation discrimination. When two sets of orientation stimuli are presented (Figure 5a), population responses for each set can be simulated and the peak of each population response can be compared to each other to simulate a trial of mean orientation discrimination (Figure 5b). This procedure was repeated for 3,000 times to make the distribution of population response differences (Figure 5c). Finally, Figure 5d shows how we constructed a psychometric function based on multiple population response differences depending on the mean orientation differences of two sets. Figure 4b shows the simulated psychometric function of Cha & Chong (2018). With the same set of parameters, Figure 4c depicts the simulated JND as a function of orientation variances of both stimuli and set size.

Psychometric functions start with a plateau that turns into a steady growth at about ∼4 degrees. This is what typically described for the variance effect on mean discrimination (Dakin, 2001; Im & Halberda, 2013; Solomon, 2010). We also observed that the greater the set size, the lower the curve which was consistent with previous studies (Dakin, 2001; Solomon, 2010; Yashiro et al., 2020). Figure 2 shows how these patterns arise from model outputs. For a fixed mean difference between two stimuli, the mean difference between the corresponding population peaks is also fixed but the SD of peak differences grows as a function of the stimulus variance. In contrast, increasing set size reduces the SD of peak differences. The mechanism of this reduction is presumably based on the *wisdom of crowds* (Alvarez, 2011; Galton, 1907), or *noise cancellation* (Baek & Chong, 2020b; Sun & Chong, 2020) that implies that averaging multiple imprecise individual estimates eventually gives more precision because uncorrelated individual errors are canceled out. Accordingly, the signal-to-noise ratio, or *population d’* decreases with increasing stimulus variance and increases with increasing set size. Thus, our model captures the pattern of data found in previous studies (Dakin 2001; Solomon, 2010; Yashiro et al., 2020). Figure 4d plots data from these studies converted to behavioral *d’* (see Figure 2 for the conversion algorithm) against the population *d’*. Pearson’s correlation between the population and behavioral *d’*s was *r* = .87 (Dakin, 2001), .67 (Solomon, 2010), and .99 (Yashiro et al., 2020) respectively.

**Figure 5.**
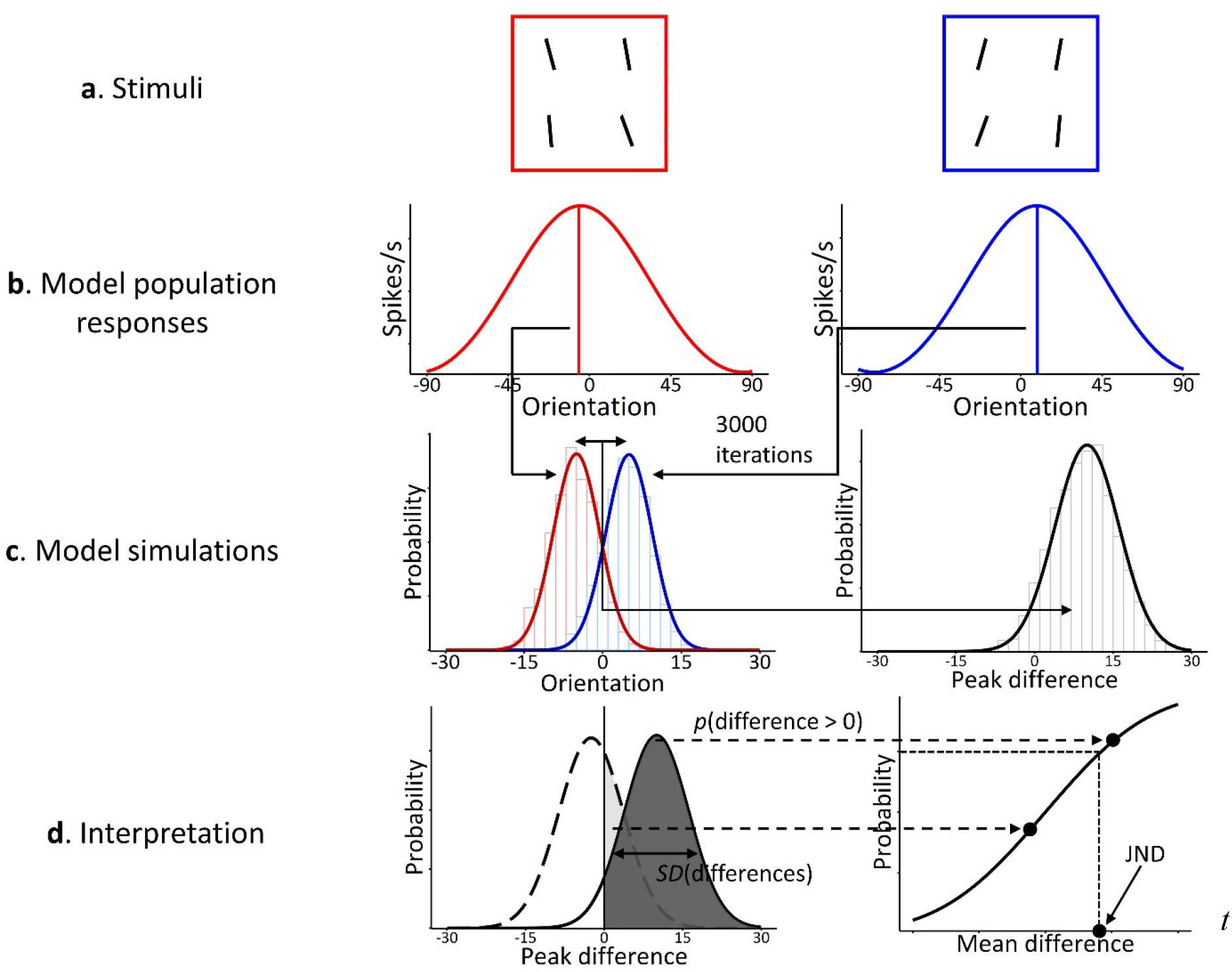
The procedure of mean orientation discrimination by the model. *Note.* (a) Two sets of oriented bars. The left set had the mean orientation of -5 deg and the right set had the mean orientation of 5 deg. (b) Model population responses to each set of stimuli. (c) We simulated population response differences for 3000 times to make their distribution for each set of stimuli with a mean difference. (d) Using these distributions, we computed the probability of having a larger population response than 0 and plotted it against mean differences. We then finally read a JND from this psychometric function.

##### Size

We also applied our model to the results of mean size discrimination (Baek & Chong, 2020a; Solomon et al., 2011). We changed only the standard deviation of tuning curve of pooling neurons to 0.3 degrees in the space where the unit was the diameter with an exponent of 1.52. We used this unit to reflect apparent sizes (Chong & Treisman, 2003; Teghtsoonian, 1965) and chose 0.21 degrees based on psychophysical findings of early noise (Baek & Chong, 2020a). We have successfully described the variance and set size effects in size averaging (Baek & Chong, 2020a; Solomon et al., 2011, Figure 6). Pearson’s correlation between the model and behavioral *d’*s was high (*r* = .98 for Baek & Chong, 2020a, and .88 for Solomon et al., 2011).

**Figure 6.**
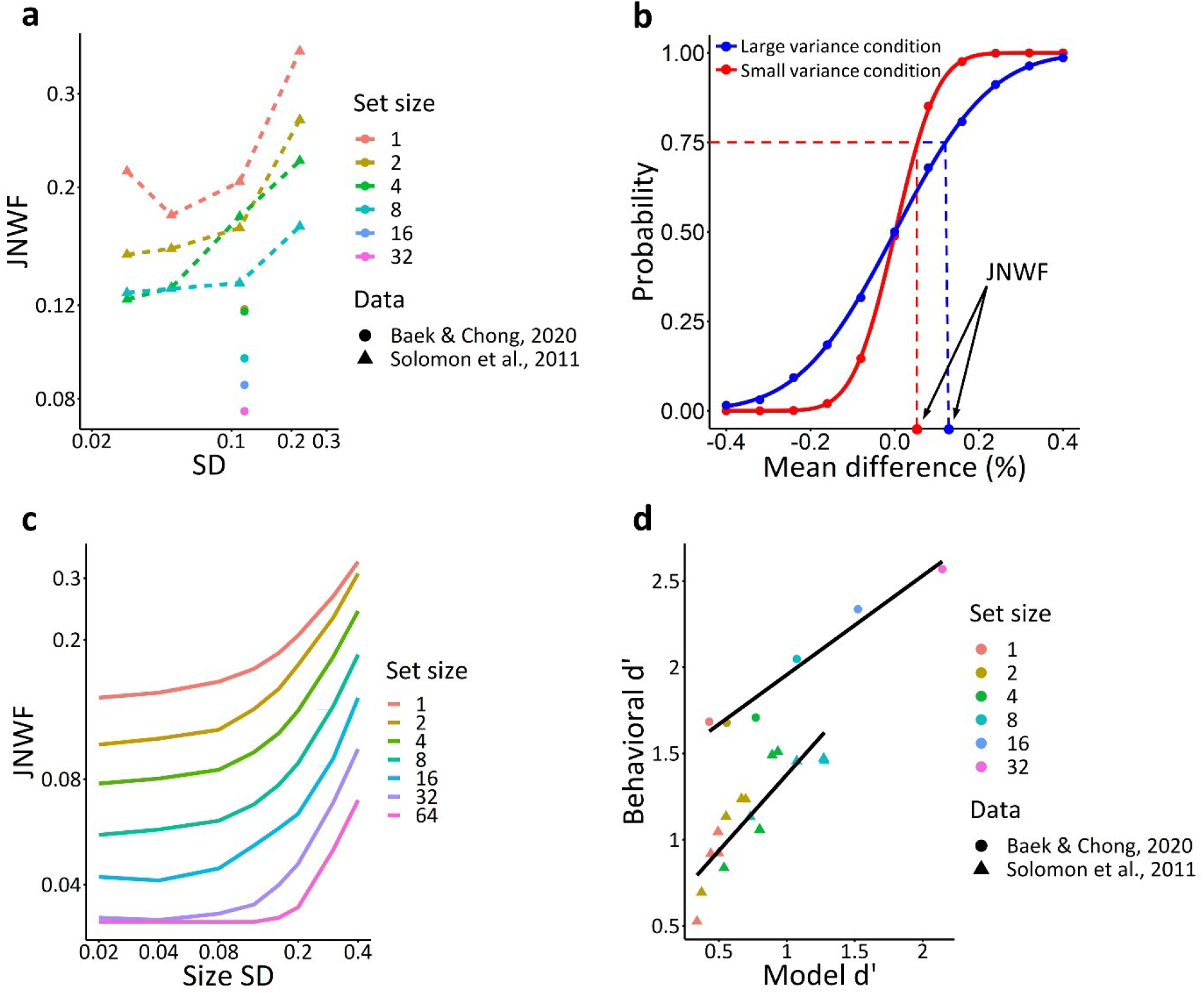
The variance and set size effects on size averaging. *Note.* (a) Previous studies have found that mean size discrimination performance decreased with increasing variance and decreasing set size (Baek & Chong, 2020; Solomon et al., 2011). (b) Psychometric functions of mean size discrimination. The blue and red lines indicate the results of the large and small variance conditions. (c) Model JNDs are plotted against SD and set size. They well capture the qualitative patterns of behavioral results presented in (a). (d) The correlation results between behavioral *d*’ from previous studies (Baek & Chong, 2020; Solomon et al., 2011) and model *d*’.

##### Motion direction and color

Here, we applied our model to both motion direction (Watamaniuk et al., 1989) and color averaging (Virtanen et al., 2020). In the case of direction averaging, we set the standard deviation of tuning curve of pooling neurons to 40 degrees with the early noise of 7 degrees (Albright, 1984; Maunsell & van Essen, 1983; Rodman & Albright, 1987). The standard deviation of tuning curve of pooling neurons was 40 degrees with the early noise of 15 degrees in the case of color averaging (Maule & Franklin, 2016). Figure 7 shows the behavioral and simulation results. The model has successfully described the results of direction averaging (Watamaniuk et al., 1989, Figure 7a) and color averaging (Virtanen et al., 2020, Figure 7b). Peason’s correlation between the model and behavioral *JND*’s was high (*r* = .98 for Watamaniuk et al., 1989, and .81 for Virtanen et al., 2020).

**Figure 7.**
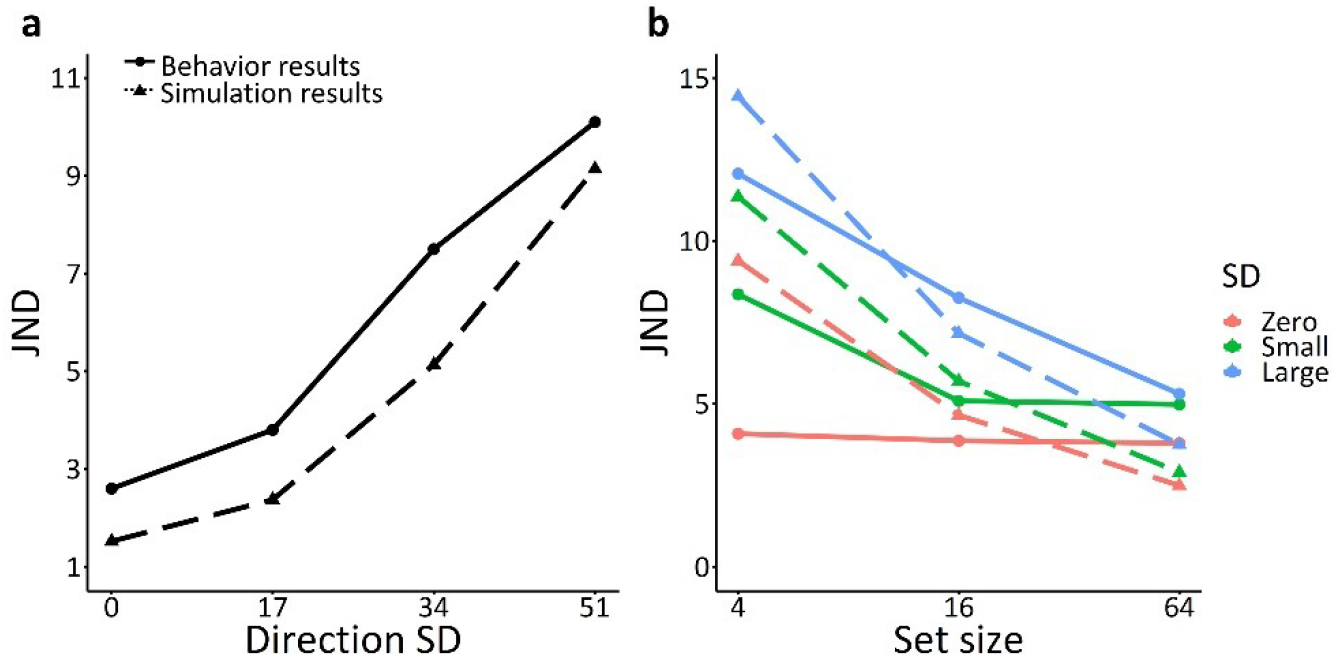
The behavior and simulation results of averaging motion directions and colors. *Note.* (a) The results of averaging motion directions (Watamaniuk et al., 1989). (b) The results of color averaging (Virtanen et al., 2020).

#### Orientation and size averaging using the adjustment method

The model can also predict the results of averaging studies using an adjustment method (Epstein et al., 2020; Khvostov & Utochkin, 2019; Kim & Chong, 2020; Maule & Franklin, 2016; Utochkin & Brady, 2020). We did not change any parameters of the model and used the peak of population response to a set of presented stimuli to predict an observer’s reported mean in each trial. We have successfully described the results of an adjustment method in orientation averaging (Utochkin & Brady, 2020, Epstein et al., 2020, Figures 8a and 8b respectively), in size averaging (Khvostov & Utochkin, 2019; Kim & Chong, 2020, Figures 8c and 8d respectively), and in color averaging (Maule & Franklin, 2016, Figure 8e). Pearson’s correlation between the predicted and reported means was high (r = .96 for Utochkin & Brady, 2020, .88 for Epstein et al., 2020, r = .55 for Khvostov & Utochkin, 2019, .78 for Kim & Chong, 2020). Maule and Franklin (2016) did not report individual observers’ results and thus we could not compute Pearson’s correlation in color averaging. Our simulation results (Figure 8e) show the similar pattern of the average observer results shown in Figure 3 of Maule and Franklin (2016). We also found that our model can predict the variance effect in an adjustment method. As the range of stimuli increased (from red to blue clouds in Figures 8a and 8c), clouds of dots dispersed more, showing the variance effect. The difference between the predicted and reported means became larger with larger variance.

**Figure 8.**
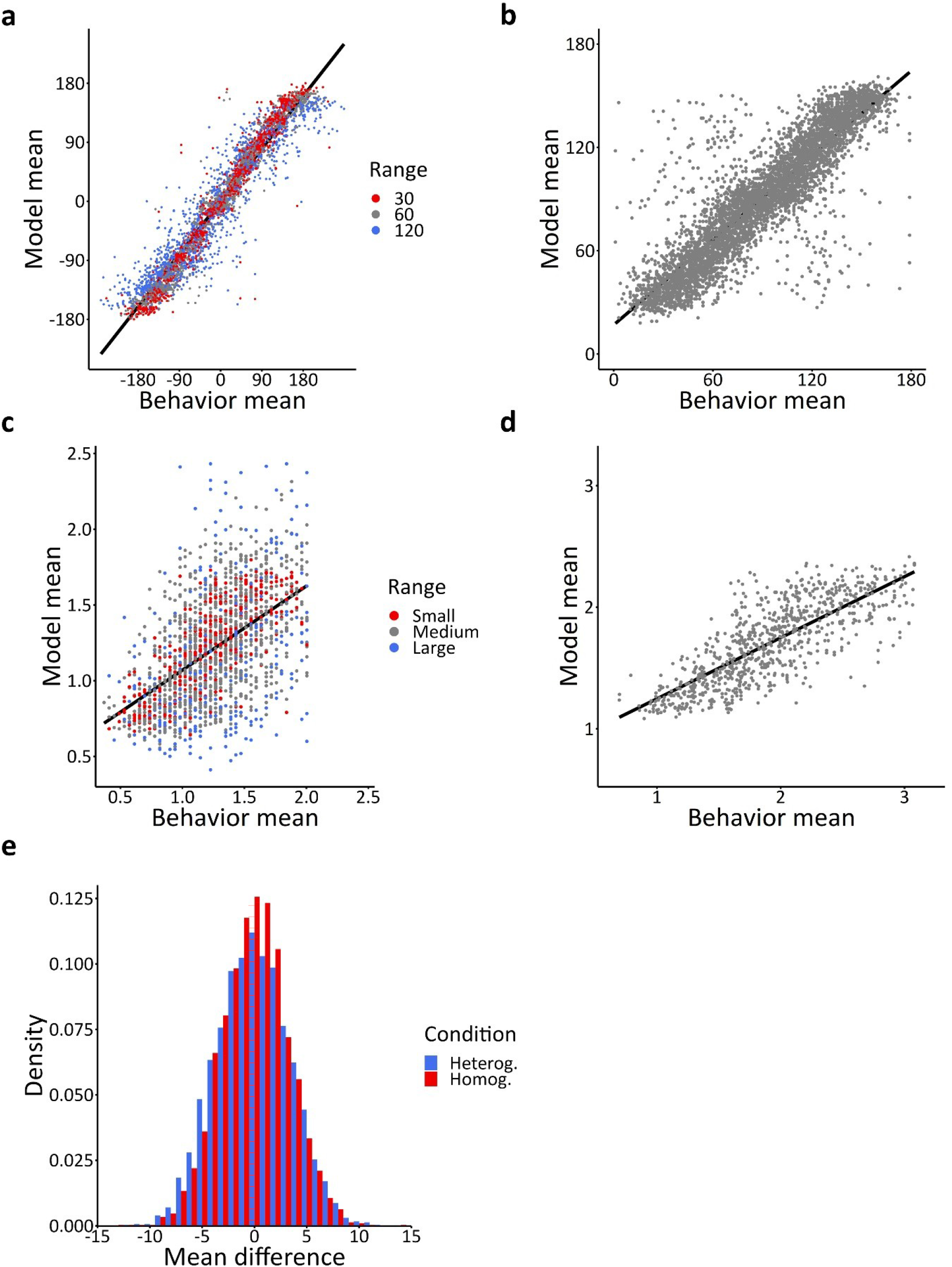
The results of an adjustment method. *Note.* Orientation averaging results (a: Utochkin & Brady, 2020; b: Epstein et al., 2020), size averaging results (c: Khvostov & Utochkin, 2019; d: Kim & Chong, 2020), and color averaging results (Maule & Franklin, 2016). Ranges in a are the same as in the original study. The range 1 in c indicates 0.3, the range 2 collapses 0.6, 0.9, and 1.2, and the range 3 indicates 1.5. We collapsed some of the conditions to visualize the variance effect better.

#### Orientation variance discrimination

Although variance discrimination is a less studied topic than averaging, several studies addressed it in the orientation domain (Jeong & Chong, 2020, 2021; Morgan et al., 2008; Norman et al., 2015). Jeong and Chong (2021) found that observers judged a display with orientations concentrated on edges (outward condition) more variable than a display with orientations concentrated on a center (inward condition, Figure 9a). Since the ranges of two displays were the same, these results suggest that people’s variance discrimination is better described by standard deviation than range. Morgan et al. (2008) and Solomon (2010) have systematically measured JND’s of variance discrimination as a function of pedestal variance (the lowest variance of the two compared sets). Morgan et al. (2008) have found a non-monotonic, “dipper” function of variance discrimination. That is, JND’s decreased with pedestal variance when it was small (up to ∼4 degrees) but the JND’s increased when the pedestal variance became larger (Figure 9b). One suggested explanation for this “dip” in the JND function is sensitivity to variance is naturally limited by the visual system’s own encoding noise. As soon as small perturbations in the appearance of individual items can equally be caused by external noise (real differences in individual orientations) or by internal noise (slight representational fluctuations of perfectly parallel individual orientations that refers mostly to early noise), the visual system cannot differentiate between these two sources of noise. Therefore, all amounts of variance not exceeding the level of internal noise are ascribed to internal and not to external noise. In other words, stimulus variances smaller than the internal noise are practically indistinguishable from each other because all real differences between them are assumed to originate from internal noise. The closer the pedestal variance approaches the limit of internal noise, the shorter the gap between this variance and the variance of another stimulus that would be able to exceed that limit and provide reliable discrimination. That is how the “dip” in the JND function arises (Morgan et al., 2008; Solomon, 2009). At the same time, Morgan et al. (2008) note that, even without noise discard explicitly defined in a model, the presence of the internal noise by itself can produce the “dip” in the JND function, although model fits in this case can be slightly worse. The “handle” part of the dipper function involves stimulus variances exceeding the amount of internal noise. Variance growth in that range quickly increases the overall noise and JND’s naturally grow with it. Note, however, that only slight tips of the “dip” in some but not all observers were found in another study of orientation variance discrimination (Solomon, 2010), as well as in the study of size variance discrimination (Solomon, 2011).

**Figure 9.**
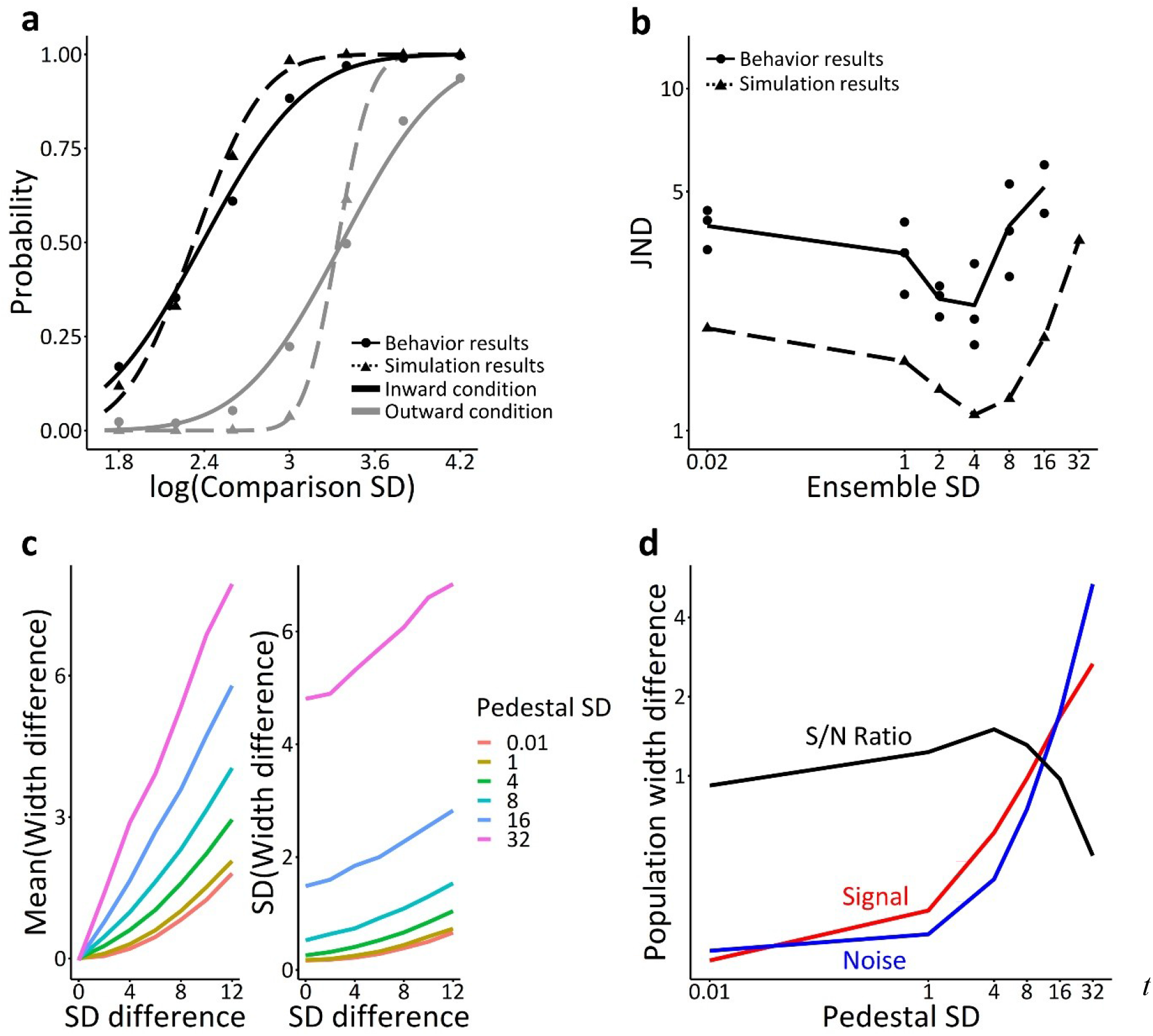
The results of orientation variance discrimination. *Note* (a) The behavioral and modeling results of Jeong and Chong (2021). Black lines indicate the inward condition and gray lines indicate outward condition. (b) The behavioral and modeling results of Morgan et al. (2008). The solid lines indicate behavioral results, and the dotted lines indicate modeling results. (c) The left panel shows how the signal strength varies depending on stimuli variance and the right panel shows how the noise changes depending on stimuli variance. (d) The signal-to-noise ratio of variance discrimination initially increases with stimuli variance and then decreases after a certain variance because of the different growth rate between the signal and noise.

Our model predicts that standard deviation is a better descriptor for people’s variance discrimination (Jeong & Chong, 2021), because standard deviation can be easily estimated by model population responses. Figure 10 depicts a psychometric function and the derivation of a JND simulated by our model that generates two population responses for each of the stimuli and compares their output bandwidths (SD’s) as proxies for variances. Figure 9a shows that psychometric functions from the model successfully simulated the inward and outward conditions from Jeong and Chong (2021). The model also explains the variance discrimination pattern as a function of physical pedestal variance (including the dip) without the need to assume a baseline amount of noise that is ascribed to the internal sources (early noise applied to each individual item independently) and discarded, as in Morgan et al. (2008). Figure 9b shows that the overall shape resembles those reported by Morgan et al. (2008), with a smooth decline before the dip and a steady increment after the dip. Figures 9c and 9d explain the emergence of the dip by the difference in relative growth of signal strength and noise in variance representation as a function of stimulus variance. It shows that the signal strength (mean bandwidth difference between two population responses) grows faster than the noise (SD of bandwidth differences) up to some point causing the signal-to-noise ratio to improve and sharpen JND’s. After that point, the signal strength grows slower than the noise and, thus, the signal-to-noise ratio steadily decreases. Note, however, that the baseline early noise in fact contributes to variance discrimination in the population coding model, as it sets the limit beyond which stimulus variance can make a meaningful change to the population width. However, no separate discard step is needed. Therefore, our model suggests a very simple mechanistic explanation of the seemingly complex, non-monotonic pattern of variance discrimination without any additional assumption. Figure 9b plots behavioral (Morgan et al., 2008) and model JNDs against ensemble set’s SD based on the model. The correlation between the behavioral and population JNDs was *r* = .71.

**Figure 10.**
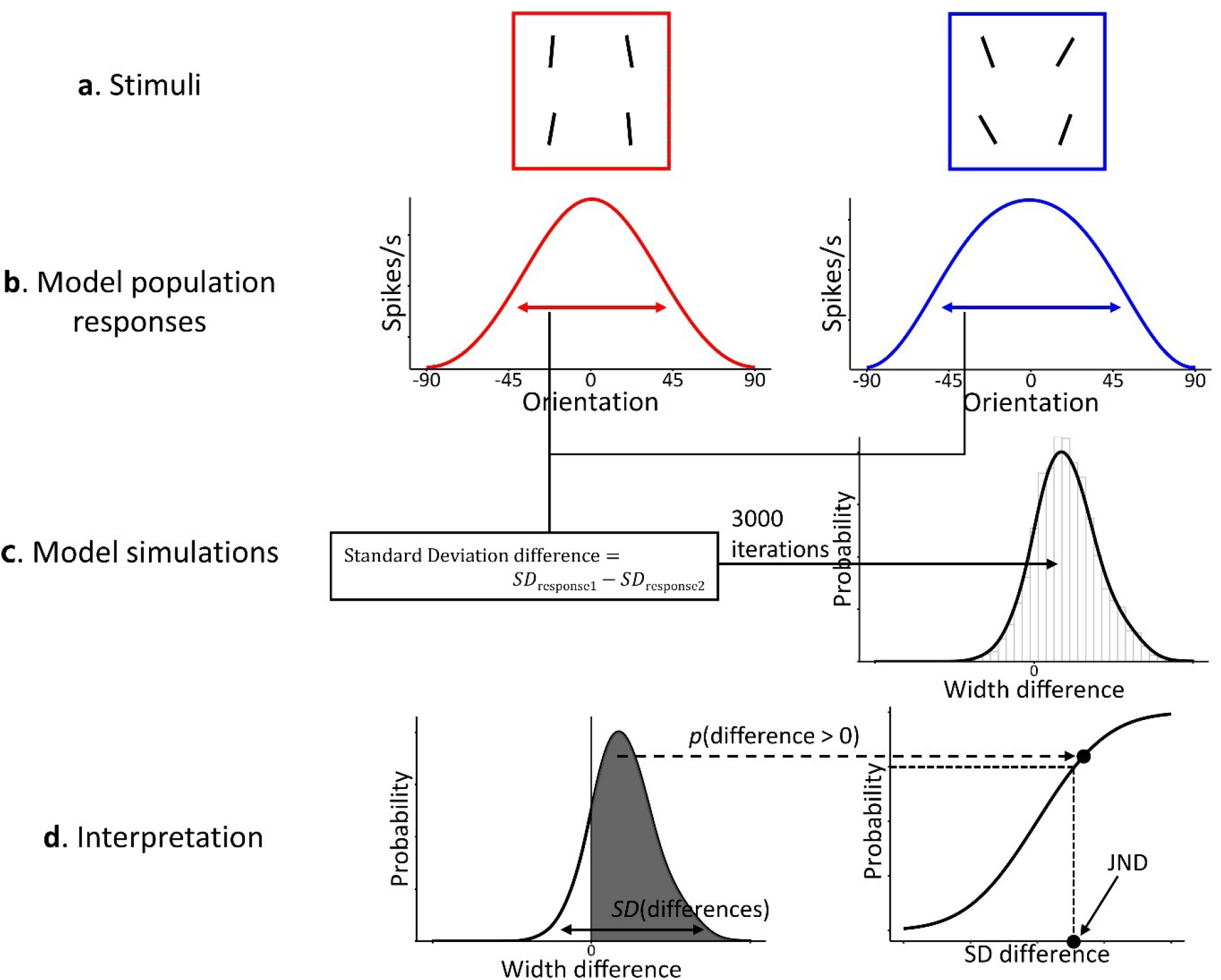
The procedure of orientation variance discrimination by the model. *Note.* (a) Two sets of oriented bars. The standard deviation of the left set was 5.8 and that of the right set was 11.5. (b) Model population responses to each set of stimuli. (c) We simulated population response differences for 3,000 times to make their distribution. (d) Based on these distributions, we constructed a psychometric function and read a JND from it.

#### Feature distribution learning

Up to this point, we applied the population coding model to various tasks requiring observers to estimate ensemble summary statistics, such as average or variance. Although these summary statistics convey the important information about all presented features, they are inevitably compressed and fail to provide more nuanced information about the representation of the whole feature distribution. However, recent research using a feature distribution learning (FDL) paradigm has shown that the visual system is able to implicitly reproduce the shape of a feature distribution, sensitive to local changes of the probability density (Chetverikov et al., 2016, 2017), which cannot be reduced to mean and variance. In the FDL task (Figure 3c), participants usually perform an odd-one-out visual search task, that is, they look for a target with a unique feature that can change unpredictably from trial to trial. Critically, the trial sequence in the FDL task is organized in streaks (e.g., 5-6 trials in a row) where distractor features are drawn from exactly the same feature distribution and switches when the distractor distribution shifts along a feature dimension and the target takes one of the values from the former distractor distribution. This allows to probe aftereffects of the previous feature distribution. Although only one point at this distribution can be probed in each streak switch, the whole distribution can be eventually constructed from multiple switches experiment-wide. One of the principal findings in the FDL research was the inhibitory effect of the former distractor distribution on the search time in switch trials. Most importantly, the amount of inhibition was proportional to the probability density of the probed feature value in the former distribution. Based on that finding, Chetverikov and colleagues (2016, 2017) concluded that the whole probability density function can be represented, yet mostly implicitly (Hansmann-Roth et al., 2021; but see Oriet & Hozempa, 2016).

Our model of ensemble population coding implies that the probabilistic feature distribution should be encoded along with summary statistics in a straightforward way. Since pooling neurons respond to input features in accordance with their tuning preferences, it is natural that more frequent features will elicit a stronger pooled response in a subpopulation of neurons tuned to these features. Thus, the overall shape of the population response necessarily reflects the physical feature distribution, as does the reaction time in the FDL paradigm. We argue that the RT functions in the FDL paradigm reflect ensemble representations in a pooled population response. This claim is based on quantitative demonstrations of strong resemblance (correlation) between the RT functions reported by Chetverikov and colleagues (2016) and population responses provided by our model. Critically, we show that this resemblance is considerably stronger than the resemblance between the RT functions and the physical feature distributions.

**Figure 11.**
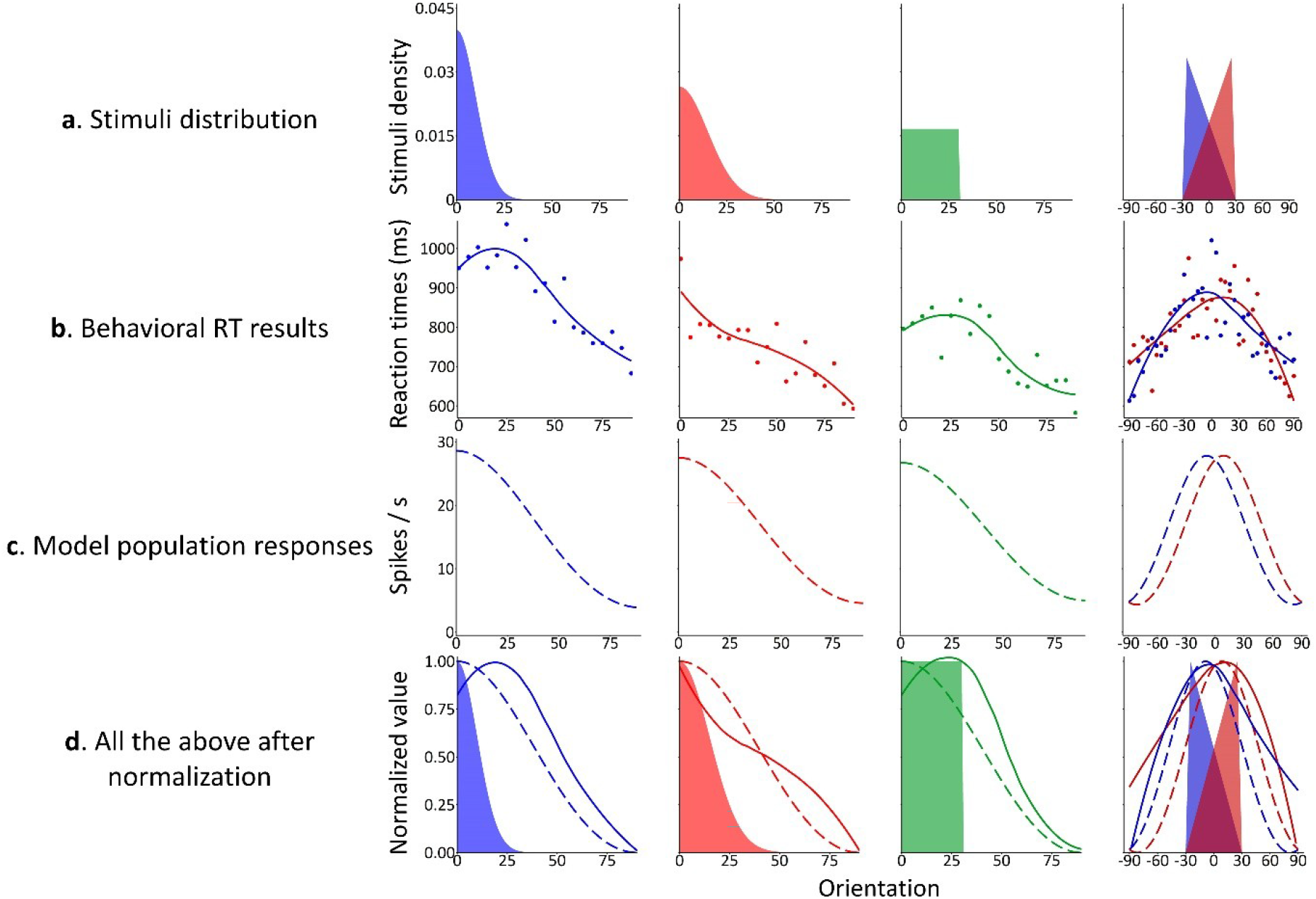
Feature distribution learning. *te.* (a) Various feature distributions used in Chetverikov et al. (2016). (b) RT results from Chetverikov et al. (2016). (c) Model population responses to each feature distribution. (d) Feature distributions, RT results, and model population responses together.

In their studies, Chetverikov and colleagues (2016, 2017) demonstrated distribution learning in orientation and color domains. The most common comparison made in these studies to demonstrate a specific effect of the distribution shape on the RT is a comparison between a Gaussian distribution and a uniform distribution (Figure 11a). Another sensitive case tested by this group is a comparison between asymmetric, triangular distributions with positive vs. negative skewness. We modeled both these cases, as they are most relevant to and illustrative of the representation of feature distribution shapes. Using the same early noise and tuning SD parameters for orientation defined in the Average discrimination in a 2-AFC task section, we found that the RT as a function of a probed orientation correlated with spike rates of model orientation-selective neurons within a range of *r* = .75 ∼ .88, which was greater than the correlations between the RT and real probability densities (*r* = .43 ∼ .69): e.g., *r* = .88 (RT-model spike rate correlation) vs. .43 (RT-real probability density correlation) in a Gaussian distribution with *SD* = 10 deg., *r* = .79 vs. .69 in a Gaussian distribution with *SD* = 15, *r* = .80 vs. .58 in a uniform distribution, *r* = .79 vs. .55 in a positively skewed triangular distribution, *r* = .75 vs. .60 in a negatively skewed triangular distribution. Note that the correlations between the RT data and the model spike rates were only slightly lower than the correlations between the RT data and regression fits to these data (*r* = .79 ∼ .93) provided by Chetverikov et al. (2016). Figure 11 shows visualizations of (a) stimuli distributions, (b) behavioral RT results, (c) model population response shapes, and (d) all three of them together that give intuitive understanding of why the population response is correlated with the RT stronger than with the actual probability density function. One strong driver of this is that the RT distributions and population responses both have extended smooth tails that the real distributions do not have and that cover a substantial range of the feature space. Second, the population response smoothens the difference between the shapes of various distributions and increases the representation at the center. This tendency is especially obvious in the comparison of two differently skewed triangular distributions where the differences between peak locations in the data and in the population codes (24 and 18 degrees, respectively) are much smaller than the difference (50 degrees) between peaks of the real distributions. This smoothing tendency also works for the uniform and Gaussian distributions (Figure 11), although our model exaggerates the resulting similarity of the corresponding population responses compared to the data. Both the extended tails and the smoothened contrasts between the distributions are explained by the way widely tuned neurons respond to individual features and then pooled together (see The Architecture of the Core model section for explanation).

#### Model extensions

Some previous studies found that averaging performance did not improve with increasing set sizes (Ariely, 2001; Marchant et al., 2013). To explain no effect of set size, we changed the amount of early noise (Figure 12a). In Figure 12a, orientation discrimination JND of 6.5 (a red dotted line) is similar across different set sizes of 2, 4, 8, if early noise increases with set sizes. Another important finding in the effect of set size on averaging is the relationship between set size and effective set size. Whitney and Yamanashi Leib (2018; see also Dakin, 2001) suggested that observers effectively used the square root of set size for averaging based on the reviews of the previous studies. To accommodate this trend, we applied late noise to our model (Figure 12b). Late noise in our model was the uncertainty of a peak selection from a population response in *L2*, expressed as the Gaussian noise with a certain SD. It indicates the degree of perturbations in ensemble estimation beyond the pooling mechanism described in our Core model. Figure 12b shows that our model explains the square-root trend better, as late noise increases. When we applied both early and late noises to our model (Figure 12c), we reproduced the pattern of results using an equivalent noise analysis reported in Dakin (2001). When early noise increases, the precision of individual orientation decreases, indicated by a large JND increment, especially around low external noise (i.e., small orientation SD). When late noise increases, the effect of external noise on averaging becomes stronger.

**Figure 12.**
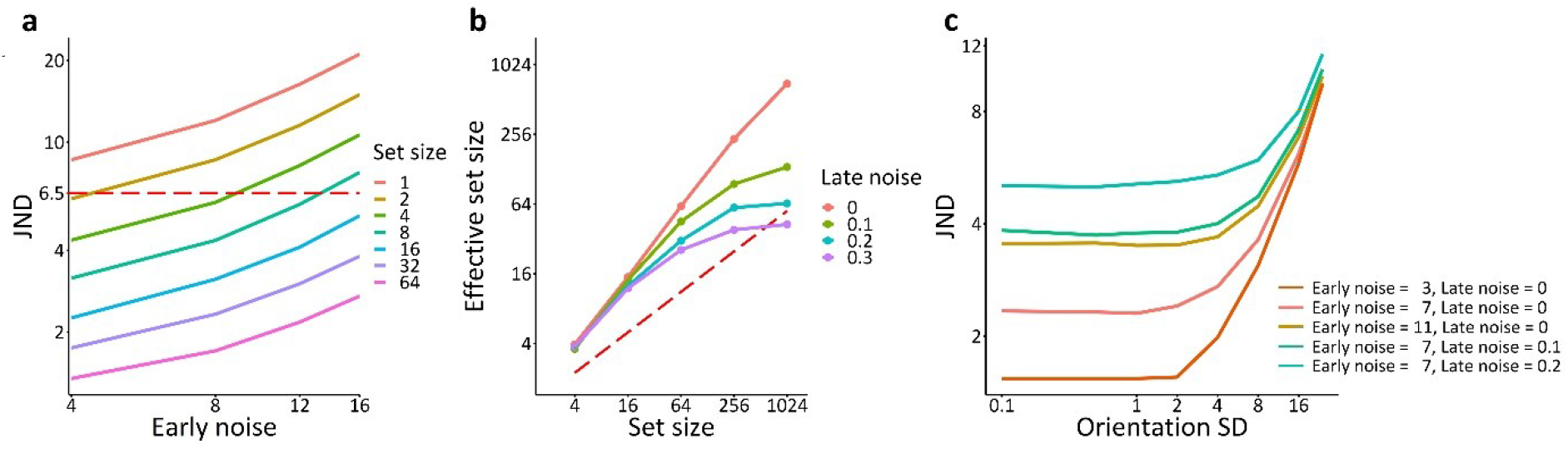
The effect of early and late noise on the population response model. *Note.* (a) Our model can explain no effect of set size on orientation averaging, if early noise increases with set sizes. Here, orientation discrimination JND of 6.5 (a red dotted line) is similar across different set sizes of 2, 4, 8, if we assume that early noise is increased with set sizes. The unit of early noise is degree. (b) Our model can also describe that the effective set size in averaging follows the square root of set size (a red dotted line), if we add late noise to the model (like the equivalent models do). The unit of late noise is spike/s. (c) With the application of early and late noise, the model can reproduce the pattern of results using an equivalent noise analysis reported in Dakin (2001).

## Discussion

The main goal of the current study was to provide a unified computational framework of ensemble perception that could account for the variety of separate findings in this field and to test it against accrued data from numerous experiments, tasks, and domains. We found that the single model of pooling and population coding captures the observed data. Specifically, the model reproduces performance in averaging tasks including most important effects such as the variability and set size effects. Furthermore, the model showed its ability to account for variance perception including the non-monotonic relationship between stimulus variance and the precision of variance estimation (the ‘dipper’ function, Morgan et al., 2008, although this effect is not always shown in data, Solomon, 2010, 2011). Since the pooled population code is a distribution of neural responses depending on a feature distribution in the stimulus, the shape of the population response in our model conveys information about the whole distribution in a natural way. This provides an account for the priming effects of the feature distribution on subsequent responses to features from that distribution (Chetverikov et al., 2017). Finally, our model provides a qualitative explanation for adaptation aftereffects of ensembles and clustering effects of similar items, which we will discuss below.

It is important to note that the model rests on the same set of assumptions and only two fixed parameters (early noise and the width of the neuron’s tuning curve) in an attempt to explain the results of previous studies across various domains and tasks, without introducing extra ad-hoc assumptions or parameters to qualitatively account for a specific effect. The late noise is an extra parameter that we added at the post-encoding stage as a model extension, and it can ultimately provide more precise quantitative fits to the data. That is, the model gives a parsimonious explanation of multiple phenomena with the same set of mechanisms. We consider our model, therefore, a neurally plausible implementation of multiple computational algorithms suggested in the previous psychophysical studies and models of ensemble perception. We would like to once again emphasize that various parameters that those models quantitatively estimate and functions that they suggest to map stimuli onto the observer’s decisions do not necessarily intend to reflect real constructs and mechanisms. A proper way of thinking about these models is as models describing algorithms, according to Marr’s (1982) classification of model levels. A failure to treat these models that way can lead to incorrect conclusions about the underlying theory of how ensemble representations are accomplished. The authors of the present article have also been trapped by the misleading interpretation of the previous observer models. Below, we will show how the concepts used by the algorithmic observer models can lead to problematic interpretations if their abstract computational characteristics are neglected. We will also show how our population coding model resolves these problems. Please note that some of the previous works explicitly used the algorithmic models of averaging to quantify observer’s inefficiency and then modeled population coding to show how the observed patterns could be implemented by the visual system (e.g., Dakin et al., 2005; Li et al., 2017).

One important example concerns the discussion of sampling efficiency in ensemble encoding. Some sampling models suggest that the effective sample size sufficient for averaging with observed levels of accuracy is a very small handful of items (e.g., Myczek & Simons, 2008). Effective sample size slowly grows with the actual set size with the rate of the square root of all presented items (e.g., Dakin, 2001; Whitney & Yamanashi Leib, 2018). This was found based on modeling and the review of previous results. If effective set size is thought of not as one of possible model parameters to quantify an observer’s inefficiency, then some serious theoretical questions can arise. For example, how does the visual system increase its capacity as a function of set size? Or why does it sample only 2 items in set size 4 when it is able to sample 4 items in set size 16? However, effective sample size may not in fact reflect a real subset of objects that observers select to estimate an ensemble summary while ignoring the rest of the objects. The pooling mechanism presented in our model does not have the effective sample size parameter and takes into account everything that falls into the receptive field of the pooling population of neurons. It turned out that, without this parameter, our model captures many crucial effects in averaging or variance estimation that are explained by effective set size in the existing algorithmic observer models.

Turning to another illustrative example, evidence has been found in some studies that, when observers judge a mean feature, they give more weight to items from the middle of the feature distribution and devalue highly deviant features (De Gardelle & Summerfield, 2011; Li et al., 2017), which makes averaging robust against the influence of unusual items and outliers (Haberman & Whitney, 2010; Epstein et al., 2020). The existing algorithmic models of this effect suggest various versions of a sampling strategy where some of the individual items gain different weights prior to averaging. For example, weights can be assigned by the likelihood estimation of each feature given the entire feature distribution. It is easy to see that the idea of likelihood estimation as a real mechanism seems to be somewhat circular or at least very complicated: It implies that the whole distribution should be known before the items are weighted and that sampling and weighting are further needed to compute the mean. In our model, robust averaging is an inherent result of the way that individual signals are pooled and decoded from the population response. The “machinery” of robust averaging is most clearly shown in our “toy” model (Figure 1), where middle neurons get more activation because they accumulate signals from both sides of the feature distribution and, thus, provide favorable conditions for peak decoding from the middle-range neurons. Indeed, as can be seen in Figure 1, activation of the middle-range neurons ‘c’ and ‘d’ on the pooled response distribution (solid black line on layer 2) is greater than activation of outlying neurons ‘a’ or ‘f’, although tuning curves for individual orientations (dashed color lines) are uniformly spanned across the entire range of neural responses. Without pooling (for example, if the mean is derived directly from the whole set of population responses to individual stimuli), decoding the mean would be less robust against outlying elements. In other previous works, it was shown that robust averaging arises from a combination of the pooled population response and a decoding rule (Iakovlev & Utochkin, 2023; Webb et al., 2007, 2010).

Importantly, our model portrays the computation of ensemble statistics as an inherent result of feedforward processing in the visual system. The mechanisms of pooling and population coding are well established in the previous research. This implies that there is no need to postulate any special “statistical processor” or any separate computational mechanism to process ensemble statistics. Rather, ensemble statistical representations can arise in the same populations of neurons that are tuned to respond to various sensory properties of stimuli. The way these neural populations represent ensemble statistics, as our model suggests, has little to do with regular mathematical computation of statistics. For example, the average ensemble feature is not accomplished by summing up individual sampled features and dividing them by a sample size, as arithmetic averaging implies. Instead, our model suggests that the average feature is accomplished more directly (without the need to represent sums and the quantities as intermediate computational steps) via the peak activation of pooled responses. Again, this sort of computation is neurally plausible, as it is in line with well established mechanisms of competitive feature interactions in large receptive fields without focused attention (Desimone & Duncan, 1995; Kastner et al., 1998; Maunsell, 2015). Although the use of arithmetic rules is based on the strict definition of what summary statistics are, it can nevertheless be misleading in terms of how it is accomplished by the visual system.

Since our model implies ensemble coding by neurons tuned specifically to visual features, it can potentially account for a set of adaptation aftereffects of ensemble statistics (Corbett et al., 2012; Jeong & Chong, 2020; Maule & Franklin, 2020; Norman et al., 2015). Adaptation aftereffects of various features are often taken as evidence for the direct coding of these features by sensory systems (Webster, 2011, 2015). Importantly, the aftereffects are usually well explained by changes in population responsiveness induced by adaptation. Since our model also has population responses in its core, it has potentially a straightforward mechanism of the ensemble aftereffects. For example, when observers are adapted to an ensemble with a large mean size and then are shown an ensemble with a smaller mean size, the peak of the population response to the latter will be shifted to the left (towards even smaller size) because the right ‘tail’ of the population response distribution overlaps with the adapted population and, hence, attenuated. This will shift the estimated mean to a smaller size – this is exactly what is observed in the data (Corbett et al., 2012). There are also adaptation aftereffects of variability that are more complicated (e.g., Jeong & Chong, 2020; Norman et al., 2015) but our model is in principle applicable to them as well, because the model also suggests the direct encoding of variability in population responses. Modeling the adaptation aftereffects of ensemble statistics is one of the future directions of developing our model.

In our model, population responses are more tightly formed with similar items. When there are two groups with sufficiently different across-group features and similar within-group features (e.g., yellowish flowers and greenish leaves), our model will form a population response with two distinct peaks and thereby distinguish two groups of features naturally without assuming additional mechanism of grouping (as shown in Figure 2a, bottom plot). This characteristic of our model can explain the difficulty of visual search, when distractors form multiple groups (Utochkin & Yurevich, 2016). It can also explain similarity-based clusters in visual working memory (Son et al., 2020). Modeling the clustering effect of ensembles is another future direction.

The idea of representing ensemble statistics via pooling in large receptive fields has repercussions in broader theories of visual perception. Ensemble statistics can be thought of as a form of gist representation (Cohen et al., 2016), that is, the rough impression of the whole scene without knowing details. A prominent ‘reverse hierarchy theory’ (Ahissar & Hochstein, 2002; Hochstein & Ahissar, 2004) links conscious perception to feedforward and feedback processing streams in the visual system. The theory suggests that the gist percept arises at the top of the fast feedforward stream, where neurons with large receptive fields respond to large portions of the visual field, whereas detailed vision requires slow feedback propagation of focused attention signals to lower-level neurons with small receptive fields. In our model, ensemble representations also emerge at neurons with large receptive fields as a result of feedforward pooling. In line with the reverse hierarchy theory, this predicts that conscious ensemble perception precedes the recognition of individual items. This prediction was recently supported neurophysiologically (Epstein & Emmanouil, 2021).

Although our model in its current simple form accounts for many basic patterns of ensemble perception, it can be further augmented with additional mechanisms to take into account other effects not covered in the current work. For example, although the current version of the model explains the core mechanism of building ensemble representations in a feedforward stream of processing, it does not consider modulatory effects of focused attention (Choi & Chong, 2020; De Fockert & Marchant, 2008; Kanaya et al., 2018; Iakovlev & Utochkin, 2021) and working memory (Williams et al., 2021) that can bias ensemble statistical estimates toward a subset of items. In our model, this can be accomplished by giving unequal weights to attended and unattended item representations when they enter or re-enter after top-down modulation from the bottom layer to the top, pooling layer. Alternatively, tuning curves of pooling neurons can be directly scaled up for attended features or scaled down for unattended features, which would reflect a neurally plausible mechanism of the biasing role of attention in neural representations within large receptive fields (Desimone & Duncan, 1995; Kastner et al., 1998; Maunsell, 2015). In addition, feedback connections from the top layer to the bottom layer are also possible. These connections can provide the modulation of low-level neural responses given the population response on top. The feedback modulation was earlier suggested as a potential mechanism of dealing with outliers (Haberman & Whitney, 2012) that allows the visual system to detect odd-one-out elements in the ensemble efficiently as a function of their difference from the rest of items (Hochstein et al., 2018; Rosenholtz, 2001). On the other hand, this feedback modulation can reduce the influence of outliers on the overall representation of an ensemble (Haberman & Whitney, 2010) in an iterative loop process (Epstein et al., 2020).

Despite its simplicity and formal appropriateness to account for ensemble perception in all domains, the model has theoretical limitations. Most substantially, our spatial pooling mechanism implies two constraints based on neural plausibility. First, there should be an ensemble (or a texture) of spatially dispersed elements in order to represent individual elements in independent receptive fields and then pool them by populations with larger receptive fields. Therefore, we do not assume that there is exactly the same mechanism for averaging temporal sequences presented in the same receptive fields (unless a temporal integrator is assumed that can pool over time). Second, the model should work for the features that can be initially processed at a level with relatively small receptive fields, so that there should be a layer with larger receptive fields that can pool from the former. It can be problematic for complex objects (e.g., multiple faces with various expressions) and scene features that are detected at very high levels of the visual cortex with very large receptive fields. One way to deal with this issue is assuming that these features can be reducible to more basic features detectable at relatively lower levels or that these basic features are represented in multivariate distributions. Alternatively, ensemble representations of high-level features can involve different mechanisms that have little overlap with low-level ensemble representations (Haberman et al., 2015).

## Concluding remarks

Ensemble representations 1) include the average information more than constituents, 2) consist of various statistical properties such as variance and distributional properties, 3) are formed over many stages of visual processing, 4) become precise if included items are similar and more items are included, and 5) are useful for many visual functions. To provide a theoretical and computational framework for these various facets of ensemble perception, we proposed a population coding model of ensemble perception. It consists of a simple feature layer and a pooling layer. We assumed ensemble representations as population responses in the pooling layer and decoded various statistical summaries from population responses. Our model successfully predicted averaging performance in orientation, size, color, and direction of motion across different tasks. Furthermore, it predicted variance discrimination performance and the priming effects of feature distributions. Finally, it explained the well-known variance and set size effects and has a potential for explaining the adaptation, clustering, and robust averaging effects.

## Footnotes

1 Note that ensemble representations can become less precise with more number of items, when included items are dissimilar (e.g., Ji & Pourtois, 2018).

2 Our model can explain no effect of set size on orientation averaging, if early noise increases with set sizes (Figure 12a).

